# Beyond beta rhythms: Aperiodic broadband power reflects Parkinson’s disease severity–a multicenter study

**DOI:** 10.1101/2025.03.11.642600

**Authors:** Moritz Gerster, Gunnar Waterstraat, Thomas S. Binns, Natasha Darcy, Christoph Wiest, Richard M. Köhler, Jojo Vanhoecke, Timothy O. West, Matthias Sure, Dmitrii Todorov, Lukasz Radzinski, Jeroen Habets, Johannes L. Busch, Lucia K. Feldmann, Patricia Krause, Katharina Faust, Gerd-Helge Schneider, Keyoumars Ashkan, Erlick Pereira, Harith Akram, Ludvic Zrinzo, Benjamin Blankertz, Arno Villringer, Huiling Tan, Jan Hirschmann, Andrea A. Kühn, Esther Florin, Alfons Schnitzler, Ashwini Oswal, Vladimir Litvak, Wolf-Julian Neumann, Gabriel Curio, Vadim Nikulin

**Affiliations:** Department of Neurology, Max Planck Institute for Human Cognitive and Brain Sciences, Leipzig, Germany; Bernstein Center for Computational Neuroscience, Berlin, Germany; Neurophysics Group, Department of Neurology, Charité – Universitätsmedizin Berlin, corporate member of Freie Universität Berlin and Humboldt-Universität zu Berlin, Berlin, Germany; Movement Disorder and Neuromodulation Unit, Department of Neurology, Charité – Universitätsmedizin Berlin, corporate member of Freie Universität Berlin and Humboldt Universität zu Berlin, Chariteplatz 1, 10117 Berlin, Germany; Einstein Center for Neurosciences Berlin, Charité – Universitätsmedizin Berlin, corporate member of Freie Universität Berlin and Humboldt Universität zu Berlin, Chariteplatz 1, 10117 Berlin, Germany; MRC Brain Networks Dynamics Unit, Nuffield Department of Clinical Neurosciences, University of Oxford, Oxford, UK; Institute of Clinical Neuroscience and Medical Psychology, Medical Faculty, Heinrich-Heine University Düsseldorf, Düsseldorf, Germany; Department of Neurosurgery, Charité - Universitätsmedizin Berlin, corporate member of Freie Universität Berlin and Humboldt-Universität zu Berlin, Berlin, Germany; Department of Neurosurgery, King’s College Hospital NHS Foundation Trust, London SE5 9RS, UK; City St George’s, University of London & St George’s University Hospitals NHS Foundation Trust, London SW17 0QT, UK; Unit of Functional Neurosurgery, Department of Clinical and Movement Neurosciences, UCL Queen Square Institute of Neurology, London, UK; Neurotechnology Group, Technische Universität Berlin, Berlin, Germany; Department of Imaging Neuroscience, UCL Queen Square Institute of Neurology, London, UK; The Biomedical Imaging Laboratory INSERM U1146 / CNRS UMR 7371 / Sorbonne University, 15 rue de l’Ecole de Médecine, 75006 Paris, France

**Author notes:** These authors contributed equally: Gabriel Curio, Vadim Nikulin.

**Keywords:** Parkinson’s disease, subthalamic nucleus, deep brain stimulation, beta oscillations, local field potentials

## Abstract

Parkinson’s disease is linked to increased beta oscillations in the subthalamic nucleus, which correlate with motor symptoms. However, findings across studies have varied. Our standardized analysis of multicenter datasets shows that small sample sizes contributed to these discrepancies—a challenge we address by pooling datasets into one large cohort (n=119). Moving beyond beta power, we disentangled spectral components reflecting distinct neural processes. Combining aperiodic offset, low beta, and low gamma oscillations explained significantly more variance in symptom severity than beta alone. Moreover, interhemispheric within-patient analyses showed that, unlike beta oscillations, aperiodic broadband power–likely reflecting spiking activity–was increased in the more affected hemisphere. These findings identify aperiodic broadband power as a potential biomarker for adaptive deep brain stimulation and provide novel insights into the relationship between subthalamic hyperactivity and motor symptoms in human Parkinson’s disease.

## Introduction

Parkinson’s disease (PD) is characterized by progressive motor impairments due to basal ganglia dysfunction. Within the basal ganglia, abnormal subthalamic nucleus (STN) activity plays a central role, exhibiting two major abnormalities in PD: excessive beta (13–30 Hz) synchronization and increased neuronal spiking activity^1,2^. Deep brain stimulation (DBS) of the STN alleviates motor symptoms and enables local field potential (LFP) recordings. Since LFPs primarily capture oscillatory activity rather than spiking, research in humans has been skewed toward studying beta oscillations.

This research focus on beta has led to numerous reports linking subthalamic beta activity with motor impairment in PD^3^. These reports inspired beta-based adaptive DBS (aDBS)—a closed-loop stimulation approach that uses beta power as an electrophysiological biomarker to adapt DBS dynamically^4^. Early trials suggest aDBS may outperform continuous DBS^5,6^, but a deeper characterization of the beta-symptom correlation may refine its clinical application. Furthermore, current aDBS implementations assume that beta power most reliably reflects motor dysfunction—a premise that remains uncertain.

While the beta-symptom correlation has been extensively studied (Table S2), its *robustness* is unclear due to methodological variability; its *replicability* awaits testing in large, diverse cohorts; and its *strength* varies considerably across studies^7–11^. Furthermore, most studies use *across*-patient correlations, though aDBS biomarkers must track symptoms *within* individuals. Finally, beta-symptom correlations are predominantly studied in the Levodopa off-state, whereas most DBS patients remain *on* medication.

To address these challenges, we conducted a multicenter STN-LFP analysis, integrating five independent datasets (Fig. 1) to create a large and heterogeneous cohort of 119 PD patients. In Part 1, we extensively characterize the beta-symptom correlation. Part 2 compares three spectral analysis frameworks to determine which best reflects neural dynamics. In Part 3, we leverage PD’s asymmetric nature and compare STN activity between more vs. less affected hemispheres, providing unique insights into how spectral features correlate with symptom lateralization at the individual patient level. We identify spectral features that correlate *within* patients and *across* medication states—two requirements for aDBS biomarkers.

**Fig. 1.**
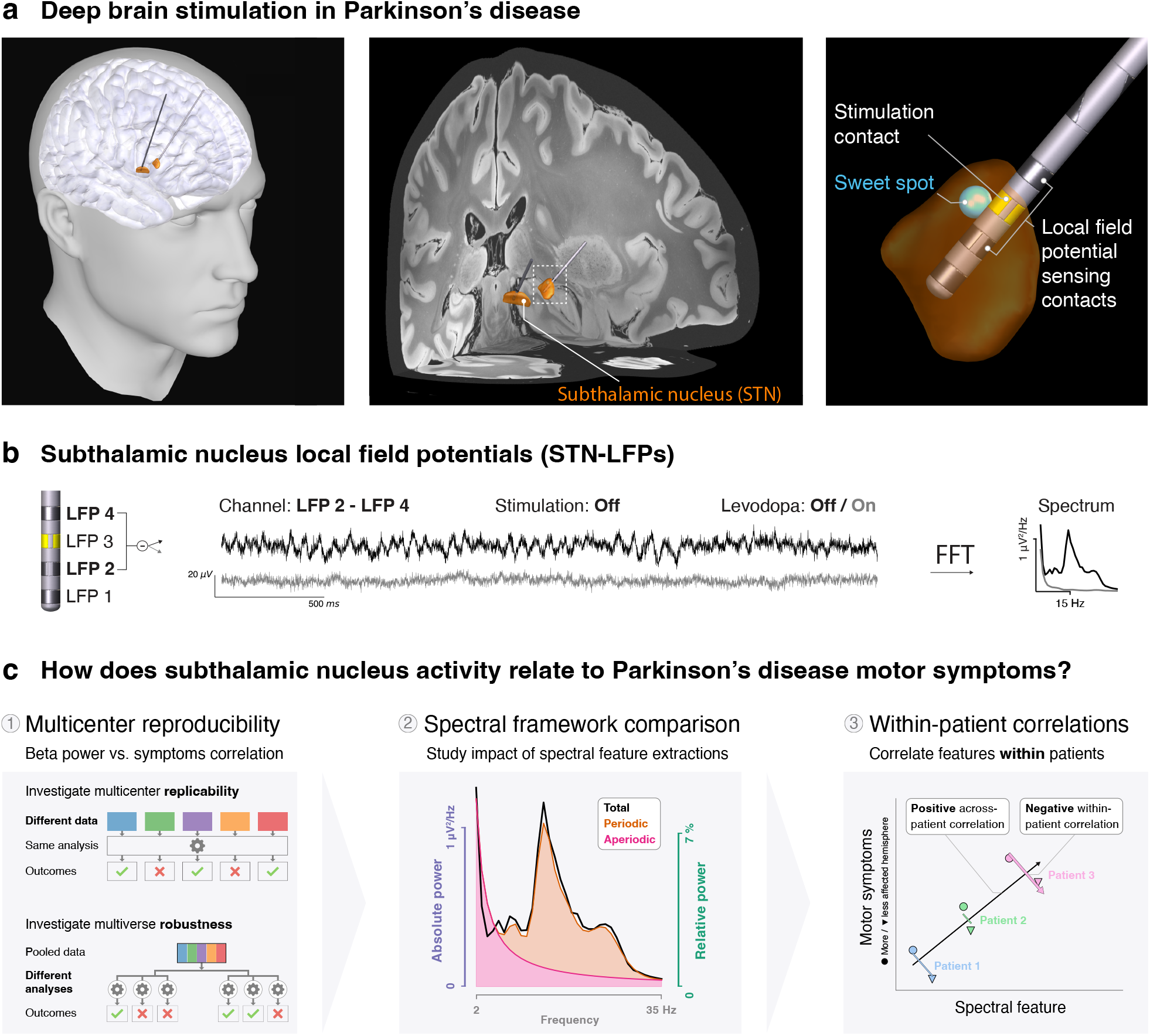
Investigating the relationship between subthalamic nucleus activity and Parkinson’s disease symptoms. **a**, Illustration of subthalamic nucleus (STN) deep brain stimulation (DBS) at three increasing anatomic scales. The 3D visualizations show a typical positioning of the DBS leads relative to the brain. The DBS contact closest to the stimulation sweet spot^12^ (blue sphere) is the estimated stimulation contact (yellow). **b**, Left: The two contacts adjacent to the stimulation contact are referenced in a bipolar montage and used for analysis (DBS leads with directional contacts are averaged so all leads have four levels of contacts). Middle: Three seconds of raw bipolar local field potential (LFP) time-series traces from one exemplary patient with and without Levodopa administration. Right: Power spectrum of the entire recording. The patient shows characteristic 14 Hz low beta oscillations off Levodopa (black), which disappear after Levodopa administration (grey). **c**, Part 1 Multicenter reproducibility: The reproducibility of the correlation between beta power and motor symptoms is assessed in terms of replicability across datasets and robustness to different analyses. Part 2 Spectral framework comparison: Different methods to extract spectral features, such as absolute total, relative total, and absolute periodic beta power, are evaluated for their motor symptom correlation. Part 3 Within-patient correlations: The relationships between spectral features and motor symptoms are tested for within-patient predictability: while within-patient correlations are imperative for successful aDBS, these might not be truthfully reflected in across-patient correlations.

We find that aperiodic broadband power meets both requirements, offering a promising target for aDBS. It is strongly elevated in the more affected hemisphere, suggesting a link to increased STN spiking activity—the second major abnormality in PD that, however, remained understudied relative to beta synchronization.

## Results

### Part 1: Multicenter reproducibility

To assess how subthalamic nucleus (STN) activity relates to Parkinson’s disease (PD) symptom severity, we analyzed resting-state STN-LFP recordings from predominantly bradykinetic-rigid PD patients. Because previous studies solely examined homogeneous single-center cohorts, we combined five published datasets to improve generalizability (Fig. 2a, Table S1, Fig. S1). Motor symptom severity was evaluated using the Unified Parkinson’s Disease Rating Scale Part 3 (UPDRS-III) with and without Levodopa medication. We calculated absolute power spectra and converted them to relative power by normalizing absolute power to the spectral sum from 5–95 Hz^13^. Fig. 2b shows an exemplary spectrum for relative (i.e., normalized) power in the Levodopa off and on conditions. We studied relative power in our reproducibility analysis to align with previous reports (Table S2).

**Fig. 2.**
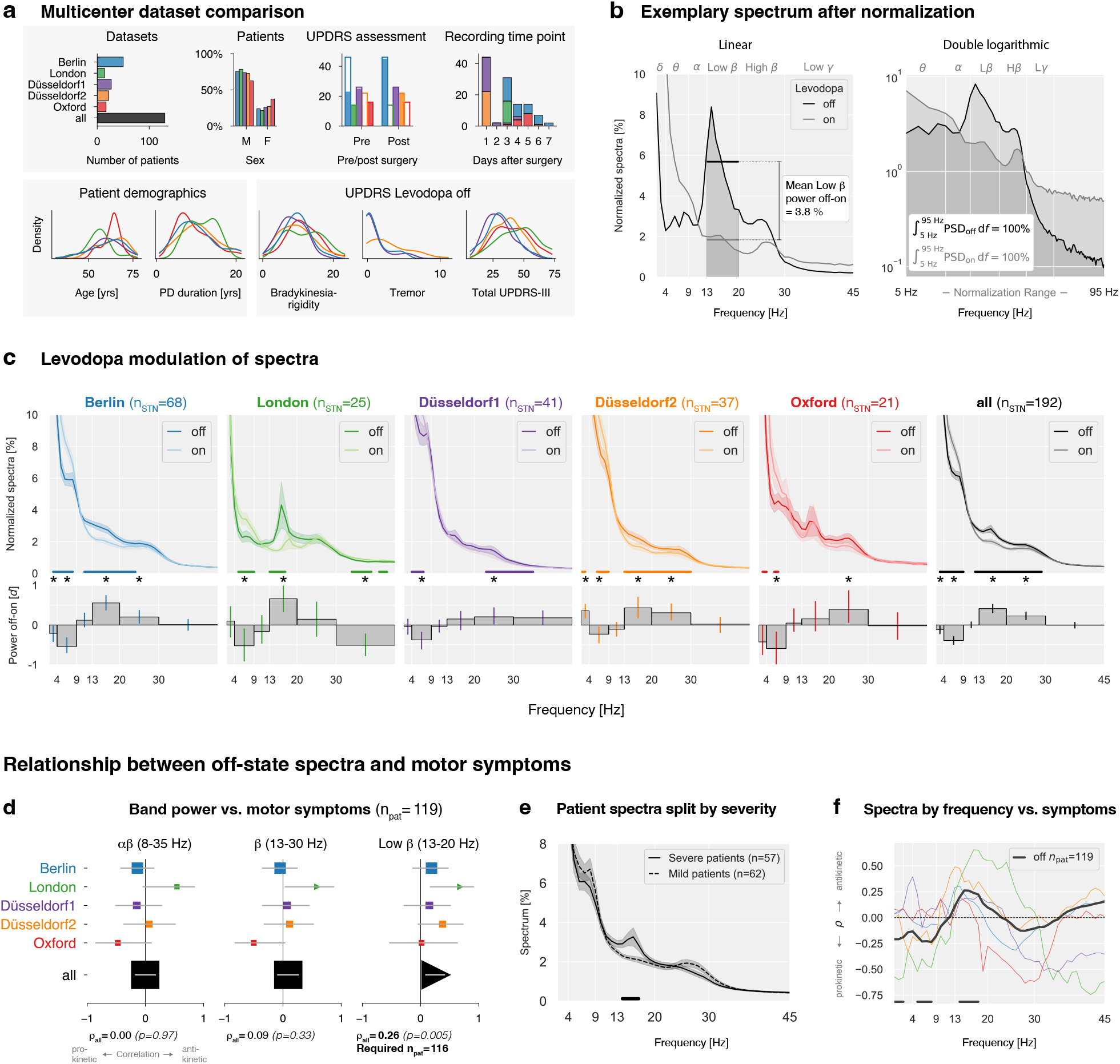
Multicenter analysis of STN-LFP recordings in Parkinson’s disease patients. **a**, Dataset and patient characteristics. Top row: Number of patients per dataset; sex distribution; time point of UPDRS assessment. Y-axis: number of patients, filled bars: proportion of evaluated patients; time point of recording. Bottom row: Kernel density estimates of age, disease duration, and UPDRS subscores, showing comparable demographic and clinical characteristics across datasets. Additional details on DBS lead manufacturers, localizations, and symptoms in the on-state are in Fig. S1. **b**, Exemplary STN-LFP power spectrum from a single subject (Berlin) in the Levodopa off and on states normalized to the 5–95 Hz frequency range. White vertical grid lines indicate frequency borders between canonical frequency bands. Left: Total relative band power was calculated as the average power within each canonical frequency band. Right: Normalization equalizes the area under the spectral curve to 100% between 5 and 95 Hz. **c**, Levodopa modulation of STN spectral power across datasets. Horizontal colored lines indicate frequency ranges with significant power differences. Below: Effect sizes (Cohen’s d) for Levodopa-induced spectral changes. Vertical colored lines indicate 95th-percentile confidence intervals; asterisks note significant effect sizes *p* < 0.05. Complete statistics in Table 1. Absolute power spectra results are in Fig. S1c. **d**, Correlations between average band power and motor symptoms. X-axis: Spearman’s correlation coefficient, y-axis: datasets, horizontal lines: 95th percentile confidence intervals, symbol sizes represent the dataset sample sizes. Triangles indicate significant correlations (*p* < 0.05); squares indicate non-significant findings. Pooled correlation coefficients and p-values are shown at the bottom. “Required npat” indicates sample size estimations (80% power requirement) for the observed correlation coefficients. **e**, Patient spectra split by median UPDRS-III score after averaging their hemispheres. The horizontal line shows a 14–17 Hz cluster of significant difference. **f**, Correlation between relative spectral power and motor symptoms for each frequency bin. Horizontal lines indicate frequencies with p-values < 0.05 for the pooled data.

### Levodopa modulation of relative power spectra

Beta-based aDBS was partly motivated by the observation that Levodopa reduces beta power^1^, but the consistency of this effect across datasets remains unknown. To assess its consistency, we examined grand mean power spectra (Fig. 2c). We used Cohen’s d to quantify Levodopa modulations, defining effect size magnitudes greater than as moderate^14^. We found at least moderate Levodopa-induced enhancements in the theta band in 4 out of 5 datasets, reductions in low beta in 3/5, and reductions in high beta in 2/5 datasets.

**Table 1.**
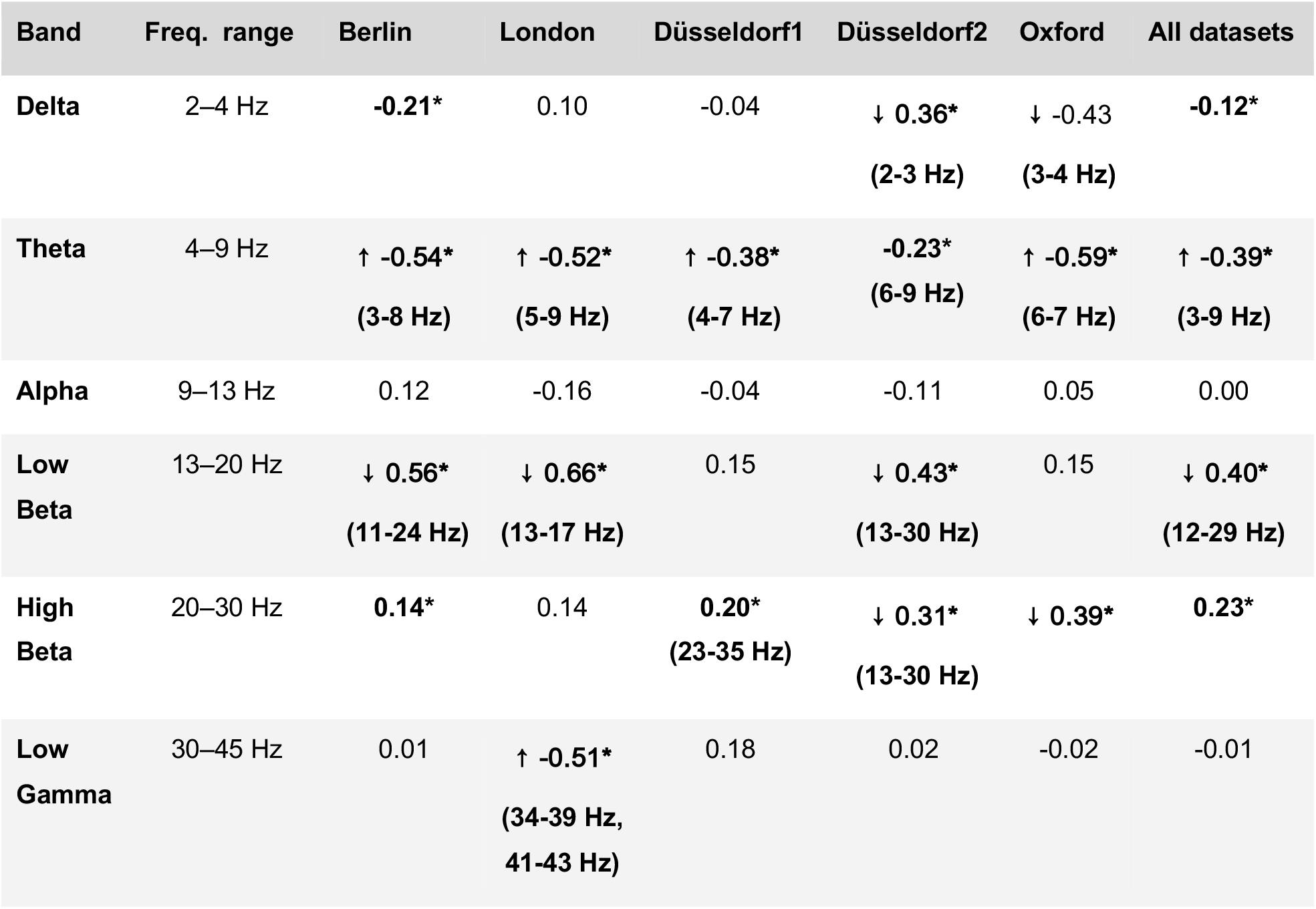
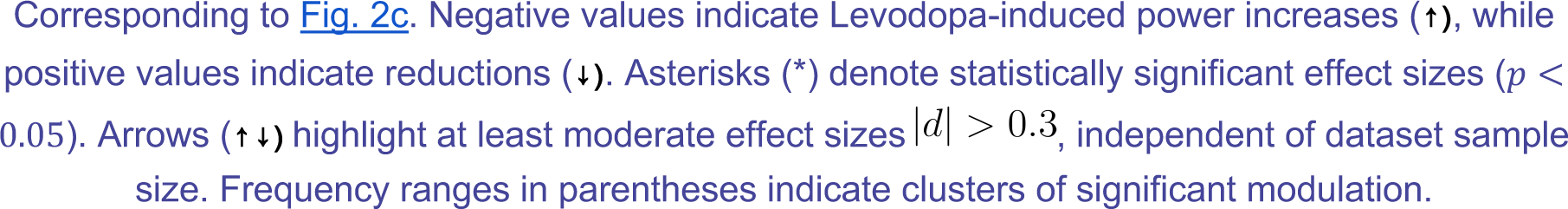
Effect of Levodopa on STN power across datasets (Cohen’s d effect sizes)

| Band | Freq. range | Berlin | London | Düsseldorf1 | Düsseldorf2 | Oxford | All datasets |
| --- | --- | --- | --- | --- | --- | --- | --- |
| Delta | 2–4 Hz | -0.21* | 0.10 | -0.04 | ↓ 0.36*<br>(2-3 Hz) | ↓ -0.43<br>(3-4 Hz) | -0.12* |
| Theta | 4–9 Hz | ↑ -0.54*<br>(3-8 Hz) | ↑ -0.52*<br>(5-9 Hz) | ↑ -0.38*<br>(4-7 Hz) | -0.23*<br>(6-9 Hz) | ↑ -0.59*<br>(6-7 Hz) | ↑ -0.39*<br>(3-9 Hz) |
| Alpha | 9–13 Hz | 0.12 | -0.16 | -0.04 | -0.11 | 0.05 | 0.00 |
| Low Beta | 13–20 Hz | ↓ 0.56*<br>(11-24 Hz) | ↓ 0.66*<br>(13-17 Hz) | 0.15 | ↓ 0.43*<br>(13-30 Hz) | 0.15 | ↓ 0.40*<br>(12-29 Hz) |
| High Beta | 20–30 Hz | 0.14* | 0.14 | 0.20*<br>(23-35 Hz) | ↓ 0.31*<br>(13-30 Hz) | ↓ 0.39* | 0.23* |
| Low Gamma | 30–45 Hz | 0.01 | ↑ -0.51*<br>(34-39 Hz,<br>41-43 Hz) | 0.18 | 0.02 | -0.02 | -0.01 |

While canonical frequency bands facilitate cross-study comparisons, they were primarily defined for cortical EEG and may not optimally capture STN oscillations. To address this, we applied non-parametric cluster-based permutation tests. When pooling all datasets, Levodopa increased power from 3–9 Hz (theta) and decreased power from 12–29 Hz (beta, Fig. 2c, Table 1). While confirming Levodopa-induced reductions in beta, we also observed strong theta enhancements. Because spectral normalization enforces interdependencies between bands, the observed theta increase may reflect a mathematical consequence of beta suppression rather than a physiological effect. To disentangle these effects, we analyze absolute power in Part 2.

#### Reproducibility of beta vs. motor symptom correlations

Although Levodopa reduces relative beta power, its pathophysiological relevance depends on its correlation with motor symptoms, which we test next.

To assess the impact of methodological choices, we conducted a multiverse analysis^15^, which tests a hypothesis across multiple analysis paths. We varied three key factors: 1) Levodopa state (off, on, or off-on difference); 2) sampling strategy (patients or hemispheres); and 3) beta frequency range. To ensure consistency with prior studies, we selected the three most commonly examined beta ranges (Fig. S2a): alpha-beta (8–35 Hz), beta (13–30 Hz), and low beta (13–20 Hz). These key factors yielded 18 distinct analyses (3 × 2 × 3).

To assess reproducibility, we examined replicability (consistency across datasets) and robustness (methodological consistency). Here, we present results for the Levodopa off-state and sampling patients (left and right STNs averaged). Results for hemisphere sampling and other Levodopa states are shown in the supplementary material (Fig. S2-S4).

Fig. 2d presents Spearman correlations between relative beta power and motor symptoms. Across datasets, the alpha-beta band did not correlate with symptom severity. Significant beta and low beta correlations emerged only in the London dataset, whereas all other datasets failed to reach significance. When pooling all datasets, only low beta power correlated with motor symptoms (ρ_*Lβ*_ *=* 0.26,*p* = 5e ^−3^).

A power analysis indicated that at least *n =* 116 patients are needed to replicate this correlation with 80% statistical power. These findings suggest that beta-symptom correlations require large cohorts (*n* > 100), precise frequency definitions (13–20 Hz), and that the relationship is weak (*r* < 0.3)^16^, prompting the question of whether additional spectral features could improve the explained variance.

#### Opposing beta and theta correlations with motor symptoms

To investigate possible spectral correlates of motor symptoms also beyond beta band frequencies, we split patients into ‘severe’ and ‘mild’ groups based on median UPDRS-III symptom scores (Fig. 2e). Patients with severe symptoms showed significantly elevated low beta power (14–17 Hz).

Next, we examined correlations between power and motor symptoms across all frequencies from 1–45 Hz using all patients (Fig. 2f). Significant negative correlations (*p* < 0.05) were found in delta (1–2 Hz) and theta (5–8 Hz), while positive correlations appeared in low beta (14–18 Hz).

Low beta correlates positively, while lower frequencies correlate negatively with motor symptoms. Because spectral normalization enforces dependencies between frequency bands, relative power may confound beta-symptom correlations and Levodopa-induced modulations. To address this, we compare spectral frameworks that normalize power, retain absolute values, or separate periodic from aperiodic components.

### Part 2: Spectral framework comparison

Spectral analysis frameworks differ in how neural power is quantified and interpreted. We compared three approaches defined by two distinctions: relative vs. absolute power and total vs. parameterized power. Relative power expresses each band as a percentage of total spectral power (5–95 Hz), while absolute power retains physical units (μV^2^/Hz).. Total power combines periodic and aperiodic components, whereas parameterization separates them, distinguishing oscillatory peaks from non-oscillatory broadband activity. We thus evaluated three frameworks: relative total power, absolute total power, and absolute parameterized power. Relative parameterized power was omitted as normalization eliminates the broadband differences that parameterization aims to model.

#### Absolute power reveals stronger Levodopa modulation of theta than beta

Before examining symptom associations, we assessed Levodopa-induced modulations using absolute power to isolate band-specific changes.

Most previous studies (25 out of 35; Table S2) analyzed *relative* power, which can complicate interpretation by conflating changes across frequency bands and masking broadband effects. For example, Fig. 3a demonstrates how absolute spectra retain Levodopa-induced broadband power modulations ((off:13.8 *μ*V^2^ vs. on:1.7 *μ*V^2^)), which are masked in relative power (Fig. 2b, same patient), as normalization equalizes the total to 100%.

**Fig. 3.**
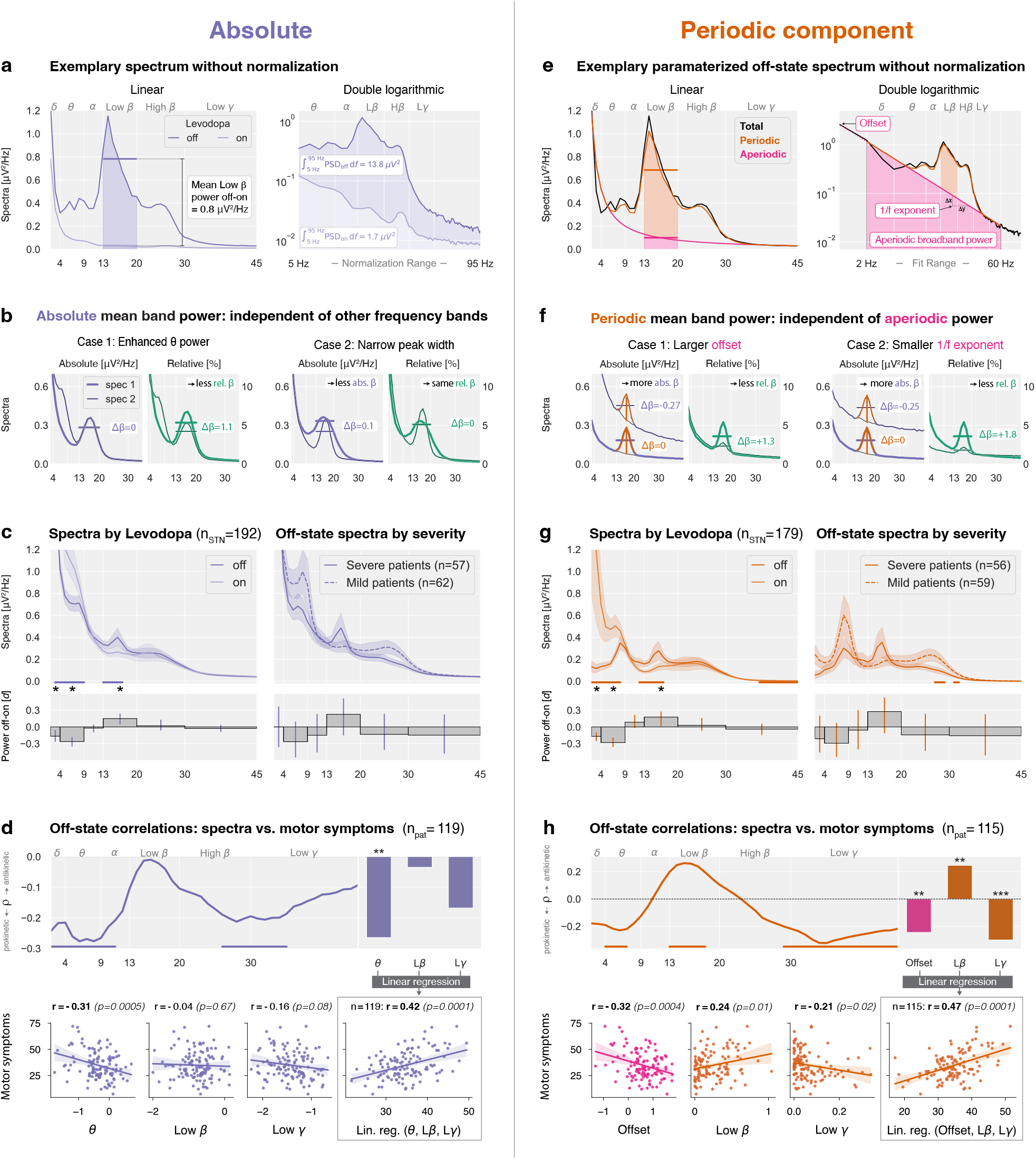
Power spectral components beyond beta are associated with symptom severity. **a**, Same spectrum as in **1b**, but without normalization. **b**, Simulations highlight the advantages of investigating non-normalized power spectra in absolute units over normalized spectra in relative units. **c**, Left: Mean absolute power spectra pooled across datasets. Bars show effect sizes, and horizontal lines indicate cluster statistics (paired statistics). Right: Levodopa off spectra for patients split by median UPDRS-III score (unpaired statistics). **d**, Top left: Correlation between motor symptoms and absolute power across frequency bins. Top right: Correlation coefficients for band power in the delta-theta, low beta, and low gamma ranges. Bottom left: Pearson correlations for linear regression features. Bottom right: Linear regression model combining theta, low beta, and low gamma power. **e**, Same spectrum as in **a** (off condition), but parameterized into periodic (orange) and aperiodic components (pink). The aperiodic component follows a 1/f power-law, modeled as where represents the offset and the exponent. **f**, Simulations highlight the implications of investigating periodic power. **g-h**, Same as **c-d** for the periodic component and aperiodic offset.

Simulations (Fig. 3b) further illustrate these distortions. An increase in absolute theta power reduces *relative* beta power, despite unchanged *absolute* beta (Case 1). Similarly, narrowing the beta peak width lowers *absolute* beta band power but leaves *relative* beta power unaffected and introduces a spurious peak increase (Case 2). These examples illustrate that absolute power more closely reflects genuine band-specific changes.

Analyzing absolute power spectra (Fig. 3c, Table 2), we observed that Levodopa significantly increased delta and theta power and decreased low beta power. Notably, high beta power remained unaffected. Effect sizes in absolute power were smaller than in relative power, where opposing modulations (theta increase vs. low beta decrease) exaggerated differences (Fig. 3b, Case 1). Crucially, absolute power revealed that Levodopa modulation was stronger in theta than in low beta frequencies (Table 2).

**Table 2.** Effect of Levodopa on STN power across spectral frameworks (Cohen’s d effect sizes)

| Band | Frequency range | Relative total | Absolute total | Absolute periodic |
| --- | --- | --- | --- | --- |
| Delta | 2–4 Hz | ↑ -0.12* | ↑ -0.17* | ↑ -0.17* |
| Theta | 4–9 Hz | ↑ -0.39*<br>(3–9 Hz) | ↑ -0.26*<br>(3–9 Hz) | ↑ -0.28*<br>(2–8 Hz) |
| Alpha | 9–13 Hz | 0.00 | -0.02 | 0.08 |
| Low Beta | 13–20 Hz | ↓ 0.40*<br>(12–29 Hz) | ↓ 0.15*<br>(13–17 Hz) | ↓ 0.18* |
| High Beta | 20–30 Hz | ↓ 0.23* | 0.02 | 0.03 |
| Low Gamma | 30–45 Hz | -0.01 | -0.03 | -0.04<br>(37–45 Hz) |
| 1/f exponent | 2–60 Hz | - | - | ↑ -0.15 |
| Offset | 2–60 Hz | - | - | ↑ -0.11 |
| Aperiodic broadband power | 2–60 Hz | - | - | 0.03 |

In OFF minus ON calculations, negative values indicate Levodopa-induced power increases (↑), while positive values indicate reductions (↓). Asterisks (\*) denote statistically significant effect sizes ( $p < 0.05$ ). Frequency ranges in parentheses indicate clusters of significant modulation.

#### Total theta power negatively correlates with symptom severity

We next examined absolute power for associations with motor symptom severity. Strongly affected patients exhibited slightly elevated low beta (*d*_*L β*_ = 0.23) and reduced theta power (*d*_*L β*_ = 0.26), though effect sizes were not significant (Fig. 3c, right). Correlation analysis (Fig. 3d) showed an overall predominance of negative correlations, suggesting a prokinetic role of broadband power, with significant negative low-frequency (1–11 Hz) and beta-gamma (26–35 Hz) correlations. The strongest correlations were observed for theta (,*r* _*θ*_ = −0.31, *p*=5e ^−4^.), while low gamma trended negatively (*r*_*Lγ*_ =−0.04,*p* = 0.08), and low beta was not significantly correlated (*r*_*Lβ*_*=* −,*p* = 0.67). A combined regression model using theta, low beta, and low gamma improved correlation strength (*r* _Lin.reg.(*θ,L, βLγ*)_ = 0.42 *p* = 1e^−4^).

#### Levodopa reduces low beta oscillations and increases theta oscillations

Conventional spectral analysis conflates periodic and aperiodic components. We applied *specparam*^*17*^ (formerly ‘FOOOF’) to separate these components (Fig. 3e). Fig. 3f illustrates the advantages of parameterizing power spectra, where a larger offset (case 1) increases absolute and decreases relative band power despite constant periodic band power. A smaller 1/f exponent has the same effect (case 2).

Analyzing absolute periodic power revealed that Levodopa significantly increased delta and theta oscillations and decreased low beta oscillations. In contrast, high beta oscillations remained unchanged (Fig. 3g). Additionally, the 1/f exponent and offset showed small but significant modulations (Table 2). These findings suggest that Levodopa-induced changes in theta and low beta power reflect true oscillatory modulations while high beta oscillations remain unaffected.

#### Aperiodic offset shows the strongest symptom correlation

To disentangle periodic and aperiodic contributions to symptom severity, we split patients based on median UPDRS-III scores (Fig. 3g, right). Low beta power was higher in severe patients (*d*_*Lβ*_ = 0.27), while theta power was lower (*d*_*θ*_ = −0.29), though neither effect reached significance. Correlation analysis (Fig. 3h) confirmed prokinetic theta (4–7 Hz), antikinetic low beta (13–18 Hz), and prokinetic low gamma (29–45 Hz) oscillations. While low beta and low gamma correlated with symptoms, the strongest association was observed for the aperiodic offset (*r*_offset_ = −0.32, *p* = 4e^−4^), which explains the negative correlation bias in absolute total power (Fig. 3d).

A combined regression model incorporating offset, low beta, and low gamma improved correlation strength ((*r* _Lin.reg.(*θ,L, βLγ*)_ = 0.42 *p* = 1e^−4^). The absolute total theta power used in the regression in Fig. 3d likely reflects both periodic theta oscillations and the aperiodic offset, given their strong correlation *r* = 0.91,*p s*= 6e^−48^ (not shown).

#### Expanding beyond beta improves correlations with symptoms

We built three linear regression models to assess how well relative total, absolute total, and absolute periodic and aperiodic power predict motor symptoms. Since relative power conflates various frequency bands, we included only total relative low beta power as a feature. To determine whether adding additional frequency bands improve symptom modeling, we incorporated theta, low beta, and low gamma power into the absolute total model (as in Fig. 3d) and the aperiodic offset, periodic low beta, and periodic low gamma power into the absolute periodic model (as in Fig. 3h). To ensure comparability, we used a matched patient set across all models (Fig. 4a), as four patients were excluded from the *specparam* analysis (Methods).

**Fig. 4.**
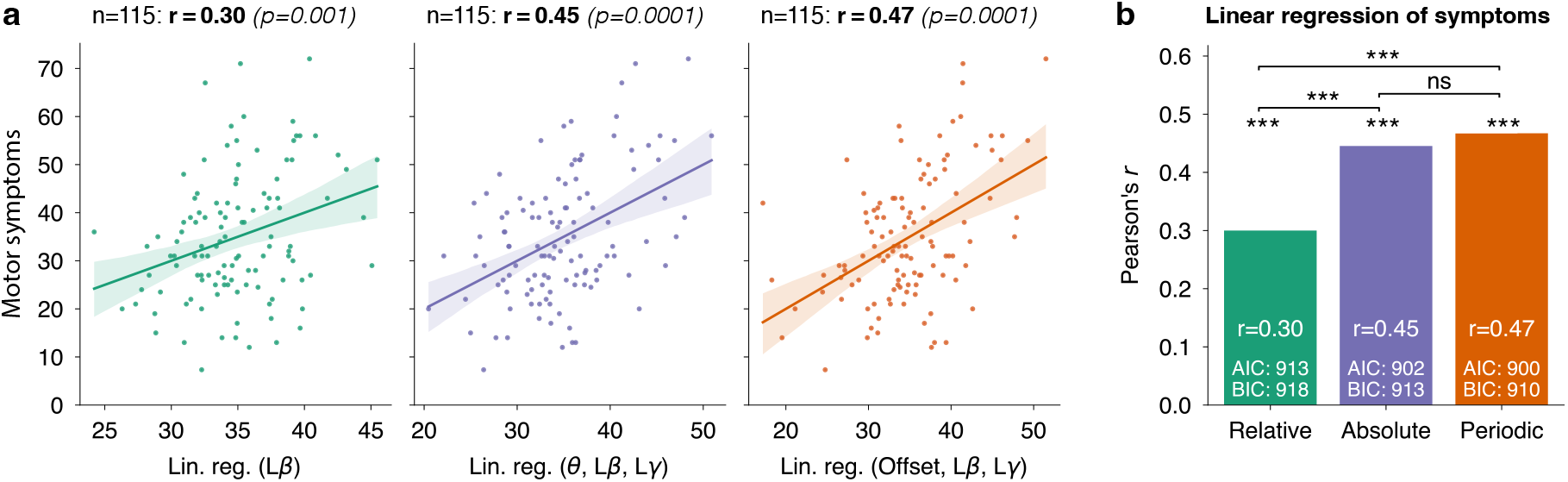
Comparison of off-state linear regression models explaining motor symptoms. **a**, Linear regression models for relative, absolute, and periodic power frameworks. The y-axis shows empirical motor symptoms (UPDRS-III scores), whereas the x-axis represents model predictions. Pearson’s correlation coefficient *r* and statistical significance *p* are shown for each model. ‘Relative’ linear regression coefficients: *b*_0_ = 27.6, *b*_*Lβ*_ = 22.3; ‘Absolute’ coefficients: *b*_*0*_ = 26.0,*b* _*θ*_ = − 14.1, *b*_*L β*_ = 12.9, *b*_*L γ*_= −10.1, ‘Parameterized’ coefficients:, *b*_0_ = 38.3,*b*_offset_ = −6.7,*b*_*Lβ*_ *=14*.*5, b*_*L γ*_= −45.1.*b*_*0*_ indicates the linear regression intercept. **b**, The absolute and periodic three-parameter models outperform the one-parameter relative model, and the periodic model outperforms the absolute model.

Performance was evaluated using-tests, alongside corrected Akaike Information Criterion (AIC) and Bayesian Information Criterion (BIC), which assess model fit while penalizing complexity. The absolute and periodic models significantly outperformed the relative low beta model (pAbs>Rel =0.0002, pPer>Rel = 2e^−5^, Fig. 4b), and the periodic model outperformed the absolute model (*p*Per>Abs = 0.03). Furthermore, AIC and BIC values were lowest for the periodic model, followed by the absolute model, and highest for the relative model.

Methodologically, these findings indicate that the periodic power framework based on absolute units best models symptom severity. Clinically, the results suggest that motor symptom severity is better explained when low beta power is complemented with low-frequency (aperiodic offset or total theta) and low gamma activity.

#### High beta oscillations co-localize with the DBS sweet spot

Beyond symptom correlations, beta power in the STN has been proposed for guiding DBS contact selection^18–23^. Here, we evaluated which spectral framework best localizes the DBS sweet spot using beta power.

We localized relative low and high beta power in the Levodopa off and on conditions (Fig. 5a). To assess spatial alignment, we correlated the localized beta power with the distance to the DBS sweet spot^12^ (blue sphere). Relative low beta power showed no significant correlation, while relative high beta power correlated negatively, indicating closer proximity to the sweet spot. These findings align with Darcy et al.^18^, who performed the same analysis in an independent cohort.

**Fig. 5.**
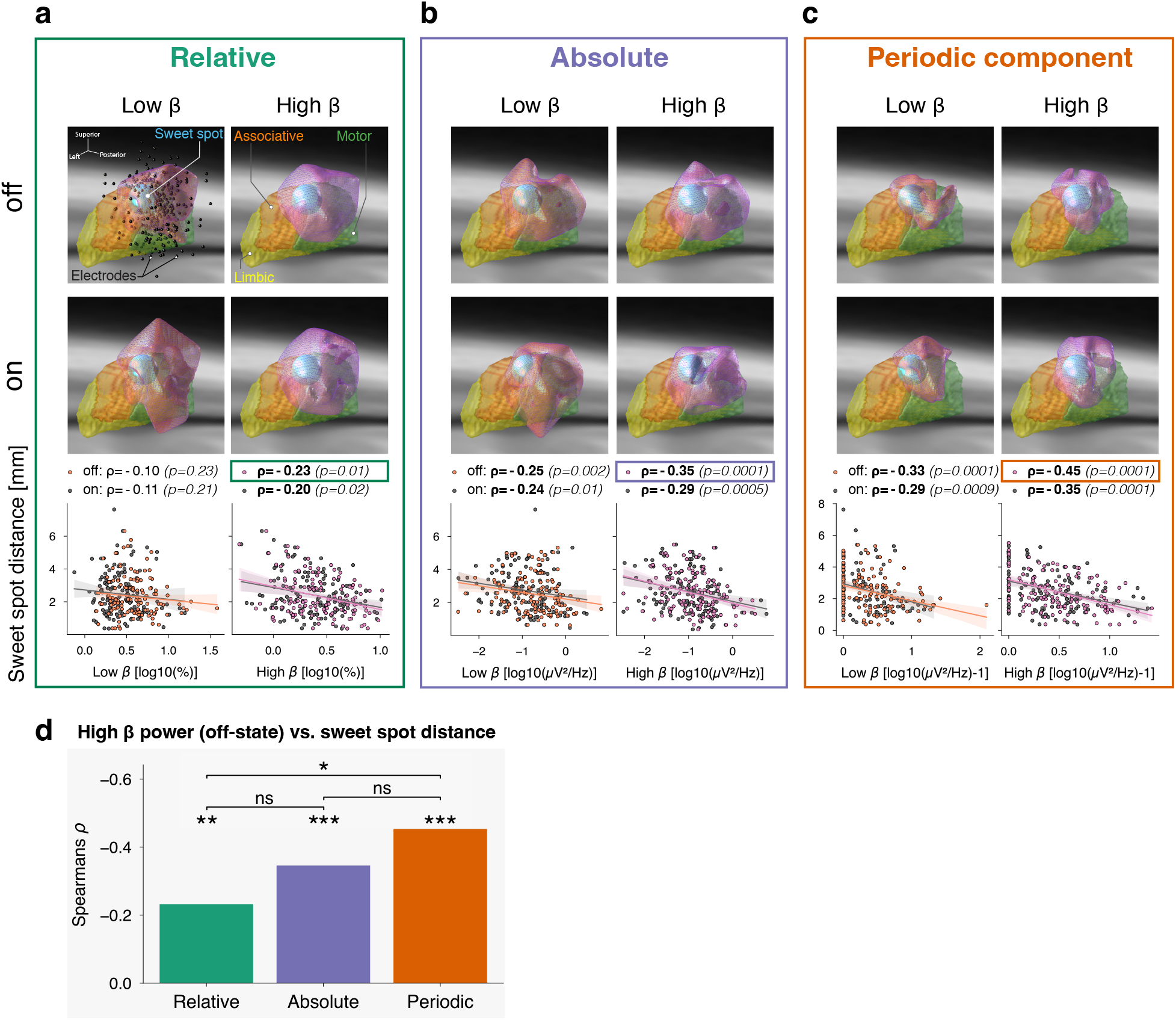
Spatial localization of beta oscillations in the subthalamic nucleus. **a**, Relative total power: Thresholded volumetric heatmaps show the spatial distribution of low beta (left) and high beta power (right) in the STN for Levodopa off (top) and on (middle) conditions. Electrode positions (black spheres) and the DBS sweet spot^12^ (blue sphere) are annotated. Bottom: Correlation between beta power and sweet spot distance, with each data point representing one STN. **b**, Absolute total power: Same as **a** without normalization. **c**, Absolute periodic power: Same as **b** after removal of the aperiodic component. High beta power in the off condition shows substantial spatial concentration at the DBS sweet spot. Sample sizes **a-c** off: n_STN_=137, on: n_STN_=132. **d**, Summary: Correlations between high beta power and sweet spot distance in the Levodopa off condition for all three spectral frameworks.

Repeating the analysis using absolute total and absolute periodic power (Fig. 5b-c) revealed consistent negative correlations across beta bands, Levodopa conditions, and both frameworks.

To compare frameworks, we performed one-tailed non-parametric permutation testing (Fig. 5d). Although absolute and periodic frameworks had comparable correlations, periodic high beta power correlated significantly more strongly with sweet spot distance than relative high-beta power (*p* =0.04). The top right panel of Fig. 5c highlights the close spatial overlap between localized periodic high beta power and the sweet spot, suggesting that beta-based DBS contact selection may be most effective using the periodic power framework.

### Part 3: Within-patient correlations

Until now, the present analyses focused on across-patient correlations in the Levodopa off-state. Critically, aDBS biomarkers must track symptoms *within* patients and *across* medication states. We treated recordings from individual hemispheres in patients as repeated measures, motivated by the asymmetry of motor symptoms typical in PD. We then inspected whether spectral features on either side correlated with lateral assessments of clinical severity.

Fig. 6a presents conceptual examples of features showing strong across-patient but weak within-patient correlations and vice versa. We included patients with recordings from both STNs and consistent symptom asymmetry across medication conditions to exclude Levodopa-induced side effects (Fig. 6b). We defined symptom asymmetry as a ≥ 1 point difference between left and right bradykinesia-rigidity scores. Relative low beta power showed no within-patient correlation (Fig. 6c).

**Fig. 6.**
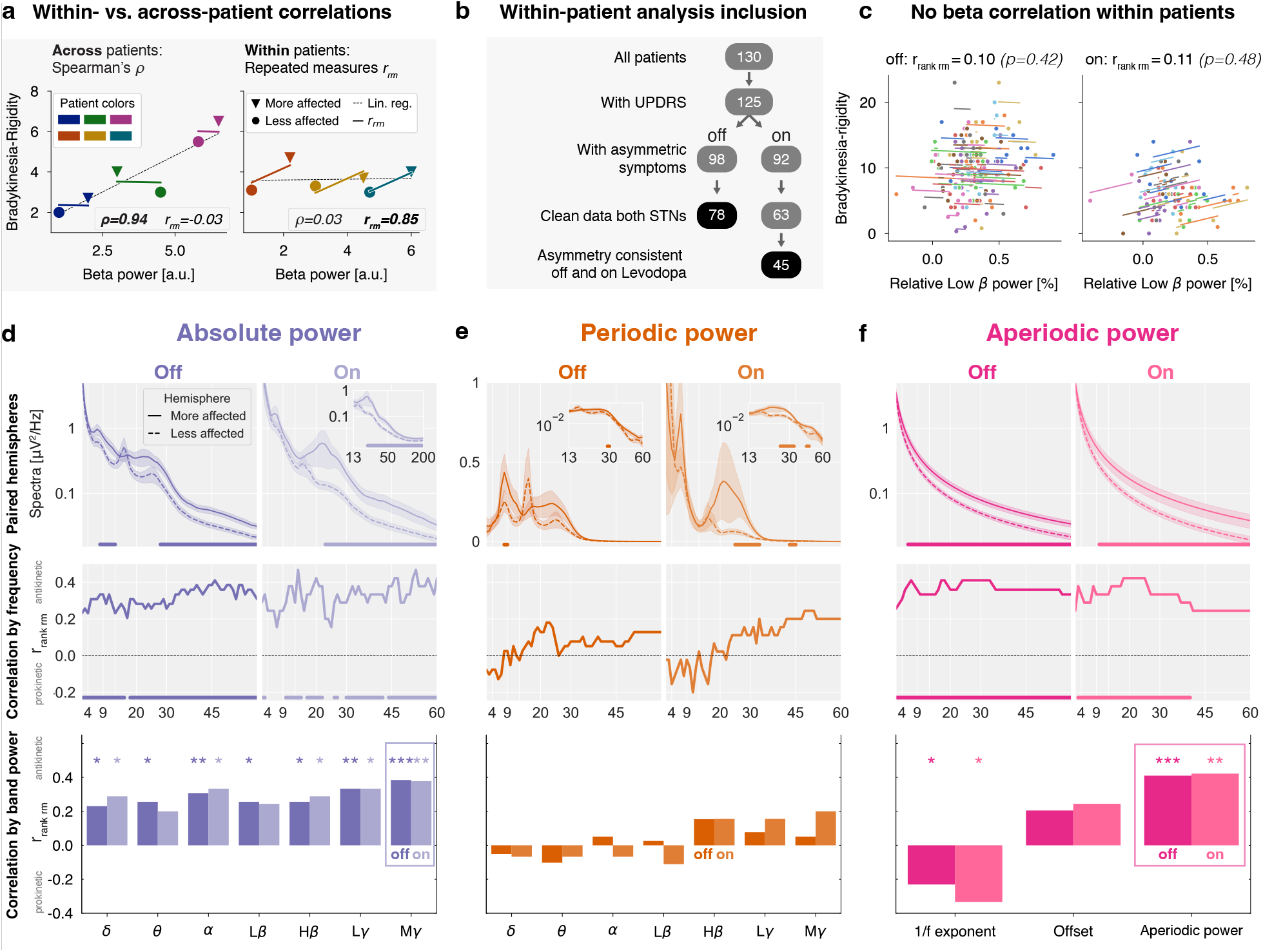
Hemispheric aperiodic broadband power correlates with symptom severity within patients. **a**, Toy example. Across-patient (left) vs. within-patient (right) correlations between beta power and bradykinesia-rigidity scores. Spearman’s rank correlation (*ρ*) quantifies across-patient relationships, while repeated measures correlation (*r*_*rm*_) quantifies within-patient relationships. Each color represents one patient, triangles and circles indicate more and less affected hemispheres. A strong across-patient correlation does not necessarily imply a strong within-patient correlation. **b**, Inclusion criteria for within-patient analysis. Black background indicates the final number of patients for each Levodopa condition. **c**, Repeated measures correlation scatter plot of relative low beta power (x-axis) and contralateral bradykinesia-rigidity score (y-axis). Each color represents a different patient, and each patient is shown with two dots indicating both hemispheres. Colored slopes indicate linear (non-ranked) repeated measures correlation. The values on top provide the repeated-measures statistics after ranking the data. **d**, Top: Absolute power within-patients paired cluster-based permutation tests for the more and less affected hemisphere in both Levodopa conditions. Middle: Repeated measures rank correlation for each frequency bin of the power spectrum and the contralateral bradykinesia-rigidity score. Bottom: Repeated measures rank correlation for each band power. Significant correlations are marked with asterisks (\**p*<0.05, \*\**p* <0.01, *** *p* < 0.00.1). **e-f**: Same as **d** for periodic and aperiodic power. Aperiodic power indicates the sum of the aperiodic component from 2 to 60 Hz.

#### Aperiodic broadband power reflects Parkinson’s disease severity

Comparing absolute spectra between hemispheres revealed elevated broadband power in the more affected hemisphere in both conditions. In the off-state, the increase was significant from 8–13 Hz and 28–60 Hz, extending to 132 Hz (not shown). The shift was significant in the on-state from 23–190 Hz (Fig. 6d, top). Separating periodic and aperiodic components demonstrated that this broadband shift primarily reflected aperiodic power (off: 6–60 Hz; on: 10–60 Hz, Fig. 6f, top), with only minor contributions from periodic oscillations (off: 8–9 Hz and 29–30 Hz; on: 25–33 Hz and 43–45 Hz, Fig. 6e, top). Normalization removed the broadband shift (Fig. S6b top).

Repeated measures correlations showed that absolute total and aperiodic power correlated with bradykinesia-rigidity scores across broad frequency ranges in both conditions. In contrast, periodic power did not (Fig. 6d-f, middle). Among canonical frequency bands, absolute mid gamma power (45–60 Hz) correlated strongest (Fig. 6d, bottom), while no periodic band correlated (Fig. 6e bottom). Aperiodic parameters showed a significant negative correlation for the 1/f exponent and a positive trend for the offset (Fig. 6f, bottom), aligning with broadband power elevation (Fig. 3f). Summing aperiodic power from 2–60 Hz, which we term ‘aperiodic broadband power’, yielded the strongest within-patient correlations (off:r_*rank rm*_ =0.41,*p* = 6e^−4^; on: r_*rank rm*_ =0.42, *p* = 0.004; Fig. 6f, bottom).

#### Total mid gamma power: A practical candidate for aDBS

An aDBS biomarker must be symptom-relevant on the individual level and readily extractable in real time. Although aperiodic broadband power strongly correlates with symptoms (Fig. 7c), its extraction requires parameterization, limiting its immediate clinical use. Instead, total mid gamma power can be rapidly obtained using simple ±2.5 Hz spectral means, as implemented in current aDBS devices (e.g., Medtronic Percept™ PC).

**Fig. 7.**
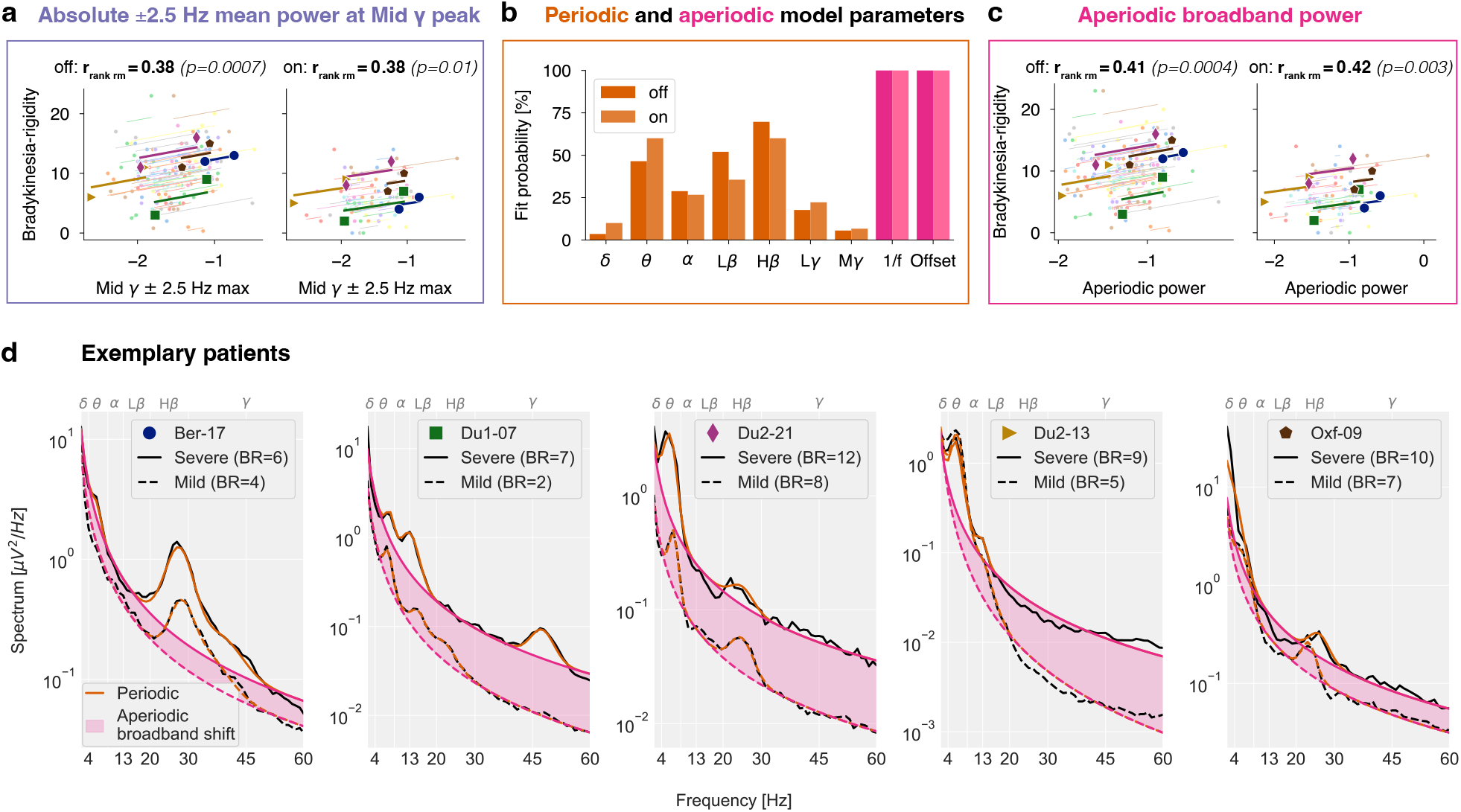
Absolute total mid gamma power as a potential adaptive deep brain stimulation (aDBS) biomarker. **a**, Mid gamma power (maximum ±2.5 Hz mean) correlates with symptom severity within patients both off and on Levodopa. **b**, Probability of detecting oscillatory peaks across frequency bands using *specparam*. **c**, Aperiodic broadband power (2–60 Hz) correlates with symptom severity in both conditions. **d**, Spectra of five representative patients in the Levodopa on condition. Symbols correspond to those in **a** and **c**. BR: Bradykinesia-rigidity subscore for each hemisphere.

Using a maximum-centered ±2.5 Hz spectral mean, absolute mid gamma power remained significantly correlated with symptoms in both conditions (off: *r*_*rank rm*_ = 0.38, *p* = 6e^−4^; on: *r*_*rank rm*_ = 0.38,*p* = 0.01; Fig. 7a). Many STNs lacked beta peaks even without medication (low beta: ~50%; high beta: ~30%; Fig. 7b), limiting their biomarker reliability. Only when considering the full alpha-beta range (8–35 Hz) did at least 88% of STNs have a peak (Fig. S6c), as previously shown^18,24,25^.

The repeated measures scatter plots (Fig. 7a, 7c) highlight five representative patients whose spectra in the Levodopa on condition consistently show elevated broadband power in the more affected hemisphere (Fig. 7d). These findings suggest that total mid gamma power captures aperiodic broadband power and has potential as an aDBS biomarker.

## Discussion

We analyzed subthalamic nucleus (STN) local field potential (LFP) recordings from Parkinson’s disease (PD) patients, emphasizing the need for large, heterogeneous cohorts to improve research generalizability. Among three spectral frameworks, parameterized absolute power spectra provided the most direct neurophysiological insights, explaining motor symptom variance best. Moreover, our findings indicate that aperiodic broadband power may serve as a novel within-patient biomarker for symptom severity.

### Part 1: Multicenter reproducibility

Reproducibility challenges in neuroscience often arise from limited statistical power^26,27^, methodological variability^28^, unpublished analysis code^29^, and cohort homogeneity^30^. While previous studies linked STN beta power to motor symptom severity, inconsistencies across reports raised the need to study robustness and replicability. Therefore, we conducted a multicenter STN-LFP comparison to evaluate these relationships across five independent datasets.

Previous studies established that Levodopa medication reduces beta power and that beta power scales with motor symptom severity. We successfully replicated these claims in a large dataset. However, significant variability across datasets challenges replicability.

Despite typical cohort sizes ranging from 14 to 50 patients, statistical outcomes varied considerably. While all datasets showed significant Levodopa-induced increases in relative theta power and four exhibited decreased relative high beta power, only three out of five datasets demonstrated significant reductions in relative low beta power.

We assessed the beta-symptom correlation using three frequency bands, two sampling strategies, and three medication states (off, on, off-on improvement), totaling 18 analyses per dataset. Given the lack of consensus in previous studies on the optimal analysis method (Table S2), we applied this multiverse approach^15^ to evaluate robustness (methodological consistency) and replicability (consistency across datasets). With five datasets, this resulted in 90 total tests (5 datasets × 18 analyses each). We found only five significant positive correlations, five significant negative correlations, and 80 nonsignificant results (Fig. 2d, Figs. S2-S4). Notably, the five positive correlations emerged from four different analysis methods, indicating that cohort differences had a stronger influence on outcomes than methodological differences.

This raises the question of whether cohort variability stems from random sampling error or systematic dataset differences. While datasets are heterogeneous regarding neurosurgery, recording setup, and patient characteristics, none displayed atypical attributes compared to prior studies (Fig. 2a, Table S1, Fig. S1). Thus, pooling all datasets aims to improve generalizability.

Pooling improved statistical power, revealing the relationship between relative beta power and symptom severity that remained obscured in single datasets. Low beta (13–20 Hz) correlated more consistently than wide-band beta (13–30 Hz) and alpha-beta (8–35 Hz) (Fig. 2d, Fig. S2-S4), supporting prior findings that clinical beta-based applications should prioritize a narrower frequency range^3,13^. These results suggest that inconsistencies in reported beta-symptom correlations primarily result from insufficient sample sizes rather than fundamental limitations of beta power as a biomarker.

We estimate that at least 116 patients are needed to replicate the correlation between relative low beta power and motor symptoms () with 80% statistical power. However, a previous large-scale study reported a lower correlation coefficient^3^, suggesting 178 patients may be required. Given that prior studies had a median sample size of only 13 patients (Table S2), these findings underscore the need for larger cohorts or within-subject analyses, such as long-term streaming data, to improve statistical power^31–33^.

### Part 2: Spectral framework comparison

In our review of prior studies (Table S2), 22 out of 35 examined relative total beta power, while only 6 assessed absolute total, 3 relative periodic, and 4 absolute periodic power. We compared these spectral analysis frameworks to determine their impact on Levodopa modulation and motor symptom correlations.

Conceptually, our simulations illustrate that relative power is difficult to interpret due to its dependency on other frequency bands. Empirically, Levodopa-induced theta power changes had a 73% larger effect size than absolute low beta power. The opposing modulations of theta and low beta inflated relative power effect sizes, with relative low beta power exhibiting a 170% larger effect size than absolute low beta power.

Our findings on relative power align with previous reports of Levodopa-induced low beta power reductions^3,18,34–36^, theta power increases^34,36–38^, and high beta power reductions^34^. However, absolute and periodic high beta power showed no Levodopa modulation, implying that the observed relative high beta power reduction is a normalization artifact, distinguishing low and high beta power^34^.

For symptom prediction, absolute power from theta, low beta, and low gamma bands outperformed relative low beta power alone. While incorporating additional bands could improve relative power models, their interpretation remains challenging due to inherent frequency conflation. In contrast, absolute power provides a simpler interpretation by isolating band-specific effects. Our reported prokinetic roles of theta and gamma have been described in PD before^39,40^; however, prokinetic subthalamic theta can also become pathological in tremor-dominant PD patients^41,42^ or dystonia^43,44^. Overall, theta, low beta, and low gamma carry crucial motor-related information, underscoring the importance of considering additional bands beyond beta to understand PD neurophysiology better.

While absolute power improves interpretability over relative power, parameterization further enhanced symptom modeling, possibly because periodic and aperiodic components reflect distinct neural processes^17^. The periodic component of low beta correlated with motor symptoms, consistent with studies isolating periodic beta oscillations in LFPs^45–47^, directly measuring neuronal beta bursting^42,43,48^, and observing worsening motor symptoms during beta-frequency DBS^49,50^.

Aperiodic offset negatively correlated with symptoms, consistent with prior work^8,46^, but its interpretation remains uncertain due to its persistent correlation with low-frequency power. While spectral parameterization provides valuable insights, determining optimal fitting parameters is challenging, particularly for STN-LFP data^51^. Thus, parameterized spectra should be interpreted cautiously, while absolute total power–omitting fitting–is more robust and compatible with real-time applications.

Despite these challenges, parameterization significantly improved a previously reported correlation between high beta power and DBS sweet spot distance^18^. Interestingly, the DBS sweet spot aligned better with high beta sources than low beta, despite its apparently stronger pathological relevance as indicated by Levodopa modulation and symptom correlation. High beta likely propagates from the motor cortex to the STN via the hyperdirect pathway^52^, inducing pathological low beta oscillations^53^. Thus, high beta power may peak where hyperdirect pathway terminals reach the STN, corresponding to the optimal stimulation site^54–57^. LFP activity could, therefore, guide DBS contact selection^58^, particularly using absolute periodic high beta power.

### Part 3: Within-patient correlations

Research on PD biomarkers in the Levodopa on-state is scarce. Moreover, a recent study emphasized the importance of distinguishing across-from within-patient correlations^33^, as the latter control for inter-subject variability–critical in heterogeneous cohorts. This motivated our search for an STN-LFP marker that tracks symptoms *within* patients *across* medication states.

Conceptually, we demonstrate that an LFP feature can correlate across but not within patients and vice versa (Fig. 6a). Empirically, this distinction is evident in our results. For example, while the aperiodic offset correlated negatively with PD symptoms across patients (off-state, Fig. 3h), it trended positively within patients (Fig. 6f). Similarly, relative and periodic low beta power correlated positively with symptoms across patients but failed to do so within patients (Fig. 6c, Fig. 6e). A prior study reported increased relative alpha-beta (8– 35 Hz) power in the more affected hemisphere for adjacent bipolar channels in the off-state^24^.

We replicated this for adjacent channels (1-2, 2-3, 3-4) but not for distant bipolar channels (1-3, 2-4, Fig. S7) used in aDBS sensing^4,59^. Furthermore, this effect was absent in the Levodopa on-state (Fig. S7), limiting its clinical applicability.

Our finding of absent within-patient beta correlations aligns with long-term streaming studies failing to predict symptoms based on beta power^7,60,61^. However, a recent multicenter clinical trial on beta-based aDBS received regulatory approval^5^, reaffirming beta’s utility as aDBS biomarker. Furthermore, STN beta activity may propagate between hemispheres, diminishing oscillatory asymmetries and affecting within-patient correlations.

In contrast to beta, absolute total broadband power (up to 200 Hz) correlated strongly with lateralized symptoms within patients, independent of medication. Spectral parameterization revealed that these broadband elevations primarily reflected aperiodic activity, characterized by larger offsets and smaller 1/f exponents. The 1/f exponent has been hypothesized to indicate excitation-inhibition balance^62^, but aperiodic activity likely reflects multiple distinct physiological processes^63^. Despite a high correlation between offsets and 1/f exponents^64^, we observed inverse correlations with symptoms. Combining these parameters as ‘aperiodic broadband power’ provided stronger within-patient symptom correlations than each parameter alone (Fig. 6f).

Why does the more affected hemisphere exhibit elevated aperiodic broadband power, and what could this signify at the neuronal level? In the cortex, neuronal spiking activity is known to increase LFP power across a wide frequency range, spanning 30–100 Hz^65–70^, up to 200 Hz^71–76^, or even the entire broadband spectrum^77,78^ —with the notable exception of the 10– 20 Hz beta range^79^. Although subcortical areas are less studied, similar trends appear in the rat hippocampus (100–600 Hz^80^), rat STN (30–100 Hz^10^), human amygdala and hippocampus (2–150 Hz^81^), and human STN in PD (55–95 Hz^82^). Applied to our findings, broadband power elevations in the more affected hemisphere may reflect increased STN spiking, consistent with evidence from PD animal models^2,83–87^ and human intraoperative microelectrode recordings^48,88,89^.

While we confirmed that Levodopa increases 1/f exponents^10,90^, aperiodic broadband power remained unchanged (Table 2), distinguishing it from beta power and suggesting they reflect independent neural processes. Moreover, while neuronal beta bursting elevates LFP beta power^91,92^, spiking outside bursts does not contribute to LFP beta power^93^, indicating that both processes are differentially reflected in the LFP. Our results suggest that beta power – likely driven by neuronal bursting–and aperiodic broadband power–likely reflecting non-burst spiking–capture distinct pathological mechanisms that differentially affect motor symptoms.

Therefore, aperiodic broadband power shows promise as a potential aDBS biomarker, possibly complementing beta. However, its extraction requires parameterization, posing challenges for real-time implementation. In contrast, absolute total mid gamma power, which captures aperiodic broadband activity, can be readily integrated into existing aDBS systems like the Medtronic Percept™ PC without additional technological advancements.

However, several limitations temper its clinical applicability. First, the correlations identified here might not be strong enough for clinical use, potentially requiring individualized machine learning models and wearable symptom-tracking devices^94^. Second, our findings stem from single time-point DBS-off resting-state recordings, leaving its dynamics during active DBS or movement unknown. Future research should assess whether absolute mid gamma power and symptoms co-fluctuate over time, especially in naturalistic settings and during DBS. More broadly, as a likely marker of neuronal spiking, aperiodic broadband power may provide valuable insights for future invasive human LFP studies, where direct spiking measurements are often unavailable.

## Methods

### Datasets

The patient demography and disease characteristics are shown in Fig. 2a, and the DBS leads and their placement are provided in Fig. S1a-c. Please refer to Table S1 and the original publications for details on the recording procedure. In brief, all Parkinson’s disease patients underwent surgery to implant DBS leads bilaterally. Recordings were performed in the days after surgery when the electrodes were still externalized. Patients were withdrawn from all Levodopa medication overnight for off-state recordings, and they received a Levodopa dose based on their usual pre-operative schedule before on-state recordings (Table S1). For the resting state recordings, patients were instructed to rest with their eyes open for at least 3 minutes. All patients provided informed consent to participate in the research, and recordings were performed according to the standards set by the Declaration of Helsinki. The study protocol for Berlin was approved by the ethics committee at Charité Universitätsmedizin Berlin, for Düsseldorf by the ethics committee of the medical faculty of Heinrich Heine University Düsseldorf, for London by the joint ethics committee of the National Hospital of Neurology and Neurosurgery and the University College London

Institute of Neurology, and for Oxford by the Health Research Authority UK, the National Research Ethics Service local Research Ethics Committee, the local ethics committee at the University of Mainz, and the South Central-Oxford C Research Ethics Committee.

### Signal processing

#### Preprocessing

All recordings were manually screened to reject bad segments and channels, then highpass filtered at 1 Hz. Before downsampling to 2000 Hz, a low-pass filter was applied to prevent aliasing.

#### Channel selection

Directional DBS contacts were averaged at each level along the electrode shaft, and we estimated the most likely monopolar stimulation contact (either contact 2 or 3) based on proximity to the DBS sweet spot^12^. When MNI coordinates were unknown, we estimated the stimulation contact based on the largest high beta power^18^. For each STN, we select one bipolar LFP for further analysis. We chose the neighboring electrodes adjacent to the estimated stimulation contact, either contacts 1-3 or 2-4, as suggested for aDBS sensing^4,59^.

#### Absolute spectra

Power spectra were computed using the Welch algorithm with a frequency resolution of 1 Hz (1-second Hamming windows) and 50% overlap between neighboring windows. Line noise artifacts were linearly interpolated in the spectrum.

#### Relative spectra

Relative power spectra in percentage units were obtained by dividing the absolute spectra by their sum from 5–95 Hz and multiplying by 100% (Fig. 2b, right).

#### Parameterized spectra

We applied *specparam*^*17*^ to separate periodic and aperiodic spectral components with the following parameters: fit range: 2–60 Hz, peak width limits: 2–12 Hz, maximum number of peaks: 4, minimum peak height: 0.1, peak threshold: 2, aperiodic mode: fixed. Fits with an value above 0.85 were kept for further processing^95^.

#### Band power

Band power was obtained from the total or periodic spectra by selecting the average power in the canonical frequency bands delta (2–4 Hz), theta (4–9 Hz), alpha (9–13 Hz), low beta (13–20 Hz), high beta (20–30 Hz), low gamma (30–45 Hz), and mid gamma (45–60 Hz).

#### Aperiodic broadband power

Aperiodic broadband power can be calculated from the offset and the *a* 1/f *m* exponent by summing the fitted aperiodic power from to :*f*_*low*_ to *f*_*high*_

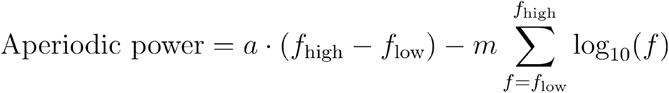

Please refer to the supplementary material for a Python implementation using *specparam*.

### Spatial localization of oscillations

The spatial localization of oscillatory activity was conducted following protocols described in Horn et al.^19^ and Darcy et al.^18^, pooling datasets from Berlin, London, and Düsseldorf1, where MNI coordinates were available. To maximize spatial resolution, we analyzed adjacent bipolar channel pairs (1–2, 2–3, 3–4), and the site of maximum band power within each STN was identified. The power was mapped into MNI space to the midpoint between bipolar recording coordinates, leading to a 4D grid (3D spatial coordinates + power values) for each frequency band and levodopa condition. Interpolations between data points were performed using a scattered interpolant, and the resulting maps were smoothed with a Gaussian kernel (FWHM = 0.7 mm)—the Lead-DBS software integrated subcortical parcellations from the DISTAL atlas^96^. Left hemisphere coordinates were flipped non-linearly to the right to increase data density and facilitate visualization. Finally, power values were thresholded at their mean plus one standard deviation to highlight the regions of interest as thresholded probability maps shown in Fig. 5.

### STN-LFP simulations

STN-LFP power spectra were simulated by constructing a Fourier power spectrum following a preset1/*f*^*m*^ power law. The corresponding phases of the Fourier spectrum are distributed uniformly randomly. To add oscillations, we add Gaussian-shaped peaks to the Fourier power spectrum with amplitudes *A* and a spectral extent given by center frequencies *f* _center_ and variances 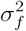. The corresponding time series, consisting of periodic oscillations and aperiodic activity, is then obtained by applying the inverse fast Fourier transform. The simulated time series have a duration of 180 s at a sampling rate of *f* _sample_ *=2400* Hz.

### Statistical analysis

Confidence intervals for all spectra were calculated through 1,000 bootstrap iterations. Differences in power spectra between the Levodopa off and on conditions were assessed using the PTE-stats toolbox (github.com/richardkoehler/pte-stats/tree/paper-moritz-gerster), employing non-parametric permutation testing with 1,000,000 permutations at an alpha level of 0.05. Cluster-based corrections were applied to adjust for multiple comparisons when significant p-values were detected, following the approach described by Maris & Oostenveld 2007^97^. Permutations were conducted by randomly reassigning conditions within subject pairs, with the mean difference used as the test statistic to compare the original and permuted datasets.

Levodopa off-on effect sizes were calculated using Cohen’s d based on the mean band power. Cohen’s d confidence intervals were calculated using 10,000 bootstrap iterations. If not indicated otherwise, correlations are computed using Spearman’s correlation. Correlation coefficient confidence intervals are calculated non-parametrically using 10,000 bootstrap iterations. Sample size estimations for correlation coefficients were computed based on Cohen 1977^16^. Within-patient correlations are visualized using repeated measures correlation^31^ and non-parametrically evaluated using ranked repeated measures correlation^32^. Direct comparison of power between more and less affected hemispheres (Fig. S7) was tested using the Wilcoxon signed-rank test.

We analyzed the relationship between STN band powers and motor symptom severity by predicting the UPDRS-III scores (−variable) using linear regression:

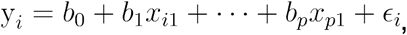

Where *i* = 1,….*n* corresponds to each patient, *b* are the regression coefficients, *p* is the number of predictors (band powers), and ϵ_*i*_ is the error term. In matrix notation, this simplifies to:

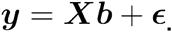

The regression coefficients were calculated by minimizing the residuals:

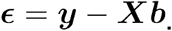

To evaluate model fits, we used Pearson’s correlation for best comparability with previous studies.

Non-nested linear regression models were compared using *J*-tests^98^. Additionally, we calculated the Akaike Information Criterion (AIC) and Bayesian Information Criterion (BIC) using the *statsmodels*.*api*.*OLS* function. To account for potential overfitting in models with small sample sizes, we applied the corrected AIC^99^

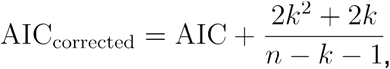

where *k* is the number of parameters.

Throughout the paper, p-values are indicated using asterisks: *p* < 0.05: ‘*’, *p* < 0. 01: ‘**’,*p* < 0.001: ‘***’.

## Data availability

The Oxford dataset is openly available at https://data.mrc.ox.ac.uk/stn-lfp-on-off-and-dbs, and the Düsseldorf2 dataset is openly available at https://openneuro.org/datasets/ds004907/versions/1.3.0 and reported in *scientific data*^*100*^. For requests regarding the Berlin dataset, please contact Wolf-Julian Neumann or the Open Data officer. For London, please contact Vladimir Litvak, and Esther Florin for Düsseldorf1.

### Code availability

All analysis code is made publicly available at github.com/moritz-gerster/STN_broadband_power. T.S.B. reviewed the analysis code for correctness.

## Acknowledgments

This work was supported by Deutsche Forschungsgemeinschaft (German Research Foundation) Project ID 424778381 TRR 295 “ReTune”. Harith Akram is supported by NIHR UCLH BRC. This work was supported by an MRC Clinician Scientist Fellowship (MR/W024810/1) held by A.O. We thank Prof. Thomas Foltynie and Prof. Patricia Limousin for their essential role in facilitating patient participation and clinical data collection at the London site. WJN received funding from the European Union (ERC, ReinforceBG, project 101077060). The 3D human head model in Fig. 1a is based on “Human Head” (https://skfb.ly/ouFsp) by VistaPrime, licensed under Creative Commons Attribution 4.0 (http://creativecommons.org/licenses/by/4.0/). We thank Dina Kemmerling for helping with the illustrations.

## Author contributions

Conceptualization, V.N., G.C.; Methodology, M.G., G.W., R.M.K, B.B., W.-J.N., G.C., V.N.; Software, M.G., T.S.B., G.W., R.M.K., N.D.; Formal Analysis, M.G.; Investigation, C.W., R.M.K., T.O.W., L.R., J.H., J.L.B., L.K.F., P.K., K.F., G.-H.S., M.S., D.T., K. A., E.P., H.A., L.Z., J.H., A.A.K., E.F., A.S., A.O., V.L., H.T., W.-J.N.; Resources, A.V., G.C., V.N.; Data Curation, M.G., T.S.B., C.W., R.M.K., J.V., T.O.W., M.S., D.T., J.H., A.A.K., E.F., A.O., V.L., H.T., W.-J.N.; Writing – Original Draft, M.G.; Writing – Review & Editing, M.G., G.W., T.S.B., R.M.K., J.V., T.O.W., J.H., L.K.F., J.H., A.A.K., E.F., A.S., A.O., V.L., W.-J.N., G.C:, V.N.; Visualisation, M.G.; Supervision, G.W., A.V., W.-J.N., G.C., V.N.; Project Administration, M.G., G.C., V.N.; Funding Acquisition, V.N., G.C., A.V.

## Supplementary Material

### Supplementary Methods

#### Dataset characteristics

**Supplementary Fig. 1.**
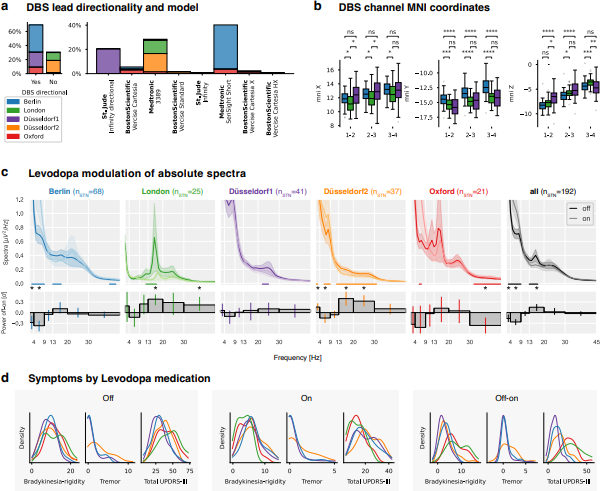
Dataset characteristics. **a**, DBS lead models by dataset. **b**, DBS channel localizations for the three datasets with available MNI coordinates. **c**, Same as Fig. 2c for absolute instead of relative power. **d**, PD symptom characteristics by medication status for all datasets.

**Supplementary Table 1.** Dataset surgery and recording details.

| Dataset | Number of patients | Number of micro recordings during surgery | Sample rate/ highpass/ lowpass | Amplifier | Patients with directional DBS leads | Recording reference | Individual On-state Levodopa dose | Off- and on-state recording on the same day | DBS center | Publications |
| --- | --- | --- | --- | --- | --- | --- | --- | --- | --- | --- |
| Berlin | 50 | 2 | 4096 Hz / - / 1600 Hz | TMSi Saga | 98% | Lowermost DBS contact (left or right) | Usual dose | No (off: day 4.3 +/- 1.3, on: 3.8 +/- 1.5) | Department of Neurosurgery at the Charité – Universitätsmedizin Berlin | 1-3 |
| London | 14 | 0 | 2400 Hz / 1 Hz / 600 Hz | CTF MEG System | 0% | Right mastoid | Usual dose | No (day 2 or 3 counterbalanced across patients) | National Hospital of Neurology and Neurosurgery and the University College London Institute of Neurology | 4 |
| Düsseldorf1 | 27 | 3.4 +/- 1.2 | 2400 Hz / - / 800 Hz | ElektaNeuro mag | 100% | mastoid | 1.5 times the Usual dose | yes | Department of Functional Neurosurgery and Stereotaxy in Düsseldorf | 5-10 |
| Düsseldorf2 | 22 | Up to 5 | 2000 Hz / 0.1 Hz / 660 Hz | ElektaNeuro mag | 0% | mastoid | 1.5 times the usual dose | yes | Department of Functional Neurosurgery and Stereotaxy in Düsseldorf | 10-12 |
| Oxford | 17 | 0 | 2048 Hz (TMSi Porti) or 4096 Hz (TMSi Saga) / - / 1600 Hz | TMSi Porti/Saga | 71% | bipolar or common average | Usual dose | yes | St. George's University Hospital NHS Foundation Trust, London (13 patients); King's College Hospital NHS Foundation Trust, London (4 patients) | 13-15 |

#### Software

Offline processing was performed with custom Python scripts using NumPy^16^, SciPy^17^, MNE-Python^18^, MNE-BIDS^19^, Pandas^20,21^, specparam^22^, seaborn^23^, Pingouin^24^, statsmodels^25^, and PTE-Stats (github.com/richardkoehler/pte-stats/tree/paper-moritz-gerster), as well as custom MATLAB scripts using SPM12^26^, and Lead-DBS^27^.

#### Aperiodic broadband power

The following code illustrates the extraction of aperiodic broadband power using *specparam*^*22*^ in the Python programming language for 2 to 60 Hz:

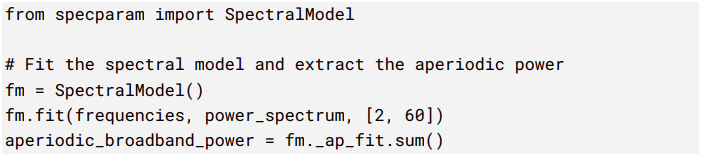

### Supplementary Results

#### Representative studies on the beta-symptom correlation

To provide context for our multicenter reproducibility analysis, we reviewed representative studies correlating macro STN-LFP resting-state DBS-off beta power with UPDRS scores. We included correlations from the baseline off-medication condition or off-on modulations of beta power and UPDRS improvements (Table S1). We found no studies specifically reporting the baseline on-medication condition. Studies calculating correlations using combined data from off and on conditions were excluded. While not exhaustive, the selected studies illustrate the variability in sample sizes (7 to 103 patients, median 13), methodologies (independent variable: absolute total, relative total, absolute periodic, or relative periodic beta power; dependent variable: lateralized UPDRS score or total UPDRS-III score), and results (17 significant vs. 22 insignificant correlations). The applied frequency borders for the “beta” band are visualized in Fig. S2a. We extracted the three most common frequency borders and referred to them as alpha-beta (8-35 Hz), beta (13-30 Hz), and low beta (13-20 Hz) bands. Studies were listed multiple times if they performed multiple analyses (such as multiple Levodopa medication conditions) to compare them with our multiverse analysis.

**Supplementary Table 2.**
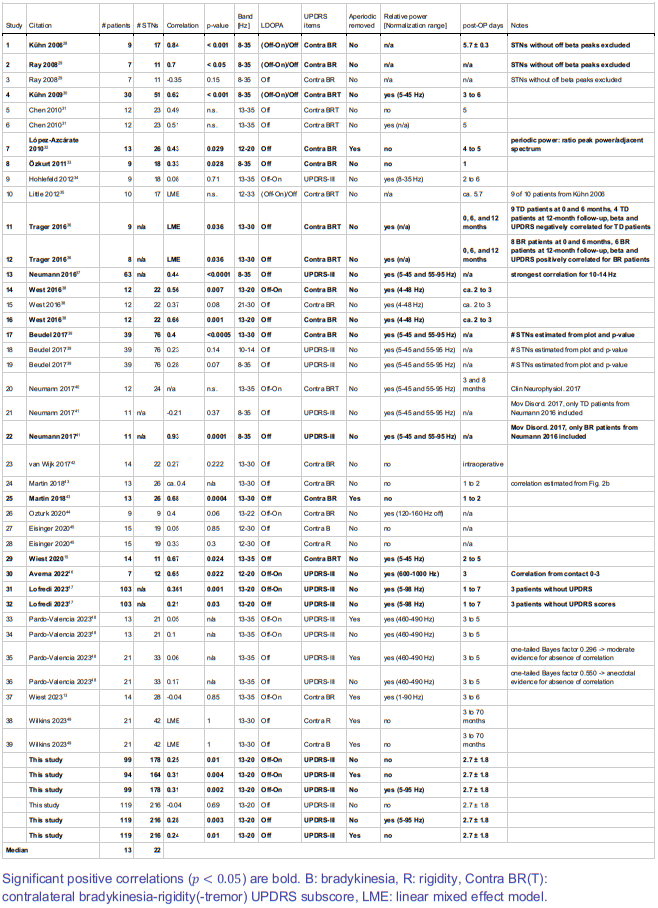
Representative selection of studies analyzing correlations between beta power and motor symptoms using macro STN-LFP recordings.

| Study | Citation | # patients | # STNs | Correlation | p-value | Band [Hz] | LDOPA | UPDRS items | Aperiodic removed | Relative power [Normalization range] | post-OP days | Notes |
| --- | --- | --- | --- | --- | --- | --- | --- | --- | --- | --- | --- | --- |
| 1 | Kühn 2006<sup>28</sup> | 9 | 17 | **0.84** | **< 0.001** | **8-35** | **(Off-On)/Off** | **Contra BR** | **No** | **n/a** | **5.7 ± 0.3** | **STNs without off beta peaks excluded** |
| 2 | Ray 2008<sup>29</sup> | 7 | 11 | **0.7** | **< 0.05** | **8-35** | **(Off-On)/Off** | **Contra BR** | **No** | **n/a** | **n/a** | **STNs without off beta peaks excluded** |
| 3 | Ray 2008<sup>29</sup> | 7 | 11 | -0.35 | 0.15 | 8-35 | Off | Contra BR | No | n/a | n/a | STNs without off beta peaks excluded |
| 4 | Kühn 2009<sup>30</sup> | 30 | 51 | **0.62** | **< 0.001** | **8-35** | **(Off-On)/Off** | **Contra BRT** | **No** | **yes (5-45 Hz)** | **3 to 6** |  |
| 5 | Chen 2010<sup>31</sup> | 12 | 23 | 0.49 | n.s. | 13-35 | Off | Contra BRT | No | no | 5 |  |
| 6 | Chen 2010<sup>31</sup> | 12 | 23 | 0.51 | n.s. | 13-35 | Off | Contra BRT | No | yes (n/a) | 5 |  |
| 7 | López-Azcárate 2010<sup>32</sup> | 13 | 26 | **0.43** | **0.029** | **12-20** | **Off** | **Contra BR** | **Yes** | **no** | **4 to 5** | **periodic power: ratio peak power/adjacent spectrum** |
| 8 | Özkurt 2011<sup>33</sup> | 9 | 18 | **0.33** | **0.028** | **8-35** | **Off** | **Contra BR** | **No** | **no** | **1** |  |
| 9 | Hohlefeld 2012<sup>34</sup> | 9 | 18 | 0.08 | 0.71 | 13-35 | Off-On | UPDRS-III | No | yes (8-35 Hz) | 2 to 6 |  |
| 10 | Little 2012<sup>35</sup> | 10 | 17 | LME | n.s. | 12-33 | (Off-On)/Off | Contra BRT | No | n/a | ca. 5.7 | 9 of 10 patients from Kühn 2006 |
| 11 | Trager 2016<sup>36</sup> | 9 | n/a | **LME** | **0.036** | **13-30** | **Off** | **Contra BRT** | **No** | **yes (n/a)** | **0, 6, and 12 months** | **9 TD patients at 0 and 6 months, 4 TD patients at 12-month follow-up, beta and UPDRS negatively correlated for TD patients** |
| 12 | Trager 2016<sup>36</sup> | 8 | n/a | **LME** | **0.036** | **13-30** | **Off** | **Contra BRT** | **No** | **yes (n/a)** | **0, 6, and 12 months** | **8 BR patients at 0 and 6 months, 6 BR patients at 12-month follow-up, beta and UPDRS positively correlated for BR patients** |
| 13 | Neumann 2016<sup>37</sup> | 63 | n/a | **0.44** | **<0.0001** | **8-35** | **Off** | **UPDRS-III** | **No** | **yes (5-45 and 55-95 Hz)** | **n/a** | **strongest correlation for 10-14 Hz** |
| 14 | West 2016<sup>38</sup> | 12 | 22 | **0.56** | **0.007** | **13-20** | **Off-On** | **Contra BR** | **No** | **yes (4-48 Hz)** | **ca. 2 to 3** |  |
| 15 | West 2016<sup>38</sup> | 12 | 22 | 0.37 | 0.08 | 21-30 | Off | Contra BR | No | yes (4-48 Hz) | ca. 2 to 3 |  |
| 16 | West 2016<sup>38</sup> | 12 | 22 | **0.66** | **0.001** | **13-20** | **Off** | **Contra BR** | **No** | **yes (4-48 Hz)** | **ca. 2 to 3** |  |
| 17 | Beudel 2017<sup>39</sup> | 39 | 76 | **0.4** | **<0.0005** | **13-30** | **Off** | **Contra BR** | **No** | **yes (5-45 and 55-95 Hz)** | **n/a** | **# STNs estimated from plot and p-value** |
| 18 | Beudel 2017<sup>39</sup> | 39 | 76 | 0.23 | 0.14 | 10-14 | Off | UPDRS-III | No | yes (5-45 and 55-95 Hz) | n/a | # STNs estimated from plot and p-value |
| 19 | Beudel 2017<sup>39</sup> | 39 | 76 | 0.28 | 0.07 | 8-35 | Off | UPDRS-III | No | yes (5-45 and 55-95 Hz) | n/a | # STNs estimated from plot and p-value |
| 20 | Neumann 2017<sup>40</sup> | 12 | 24 | n/a | n.s. | 13-35 | Off-On | Contra BRT | No | yes (5-45 and 55-95 Hz) | 3 and 8 months | Clin Neurophysiol. 2017 |
| 21 | Neumann 2017<sup>41</sup> | 11 | n/a | -0.21 | 0.37 | 8-35 | Off | UPDRS-III | No | yes (5-45 and 55-95 Hz) | n/a | Mov Disord. 2017, only TD patients from Neumann 2016 included |
| 22 | Neumann 2017<sup>41</sup> | 11 | n/a | **0.93** | **0.0001** | **8-35** | **Off** | **UPDRS-III** | **No** | **yes (5-45 and 55-95 Hz)** | **n/a** | **Mov Disord. 2017, only BR patients from Neumann 2016 included** |
| 23 | van Wijk 2017<sup>42</sup> | 14 | 22 | 0.27 | 0.222 | 13-30 | Off | Contra BR | No | no | intraoperative |  |
| 24 | Martin 2018<sup>43</sup> | 13 | 26 | ca. 0.4 | n/a | 13-30 | Off | Contra BR | No | no | 1 to 2 | correlation estimated from Fig. 2b |
| 25 | Martin 2018<sup>43</sup> | 13 | 26 | **0.68** | **0.0004** | **13-30** | **Off** | **Contra BR** | **Yes** | **no** | **1 to 2** |  |
| 26 | Ozturk 2020<sup>44</sup> | 9 | 9 | 0.4 | 0.06 | 13-22 | Off-On | Contra BR | No | yes (120-160 Hz off) | n/a |  |
| 27 | Eisinger 2020<sup>45</sup> | 15 | 19 | 0.05 | 0.85 | 12-30 | Off | Contra B | No | no | n/a |  |
| 28 | Eisinger 2020<sup>45</sup> | 15 | 19 | 0.33 | 0.3 | 12-30 | Off | Contra R | No | no | n/a |  |
| 29 | Wiest 2020<sup>15</sup> | 14 | 11 | **0.67** | **0.024** | **13-35** | **Off** | **Contra BRT** | **No** | **yes (5-45 Hz)** | **2 to 5** |  |
| 30 | Averna 2022<sup>46</sup> | 7 | 12 | **0.65** | **0.022** | **12-20** | **Off-On** | **UPDRS-III** | **No** | **yes (600-1000 Hz)** | **3** | **Correlation from contact 0-3** |
| 31 | Lofredi 2023<sup>47</sup> | 103 | n/a | **0.361** | **0.001** | **13-20** | **Off-On** | **UPDRS-III** | **No** | **yes (5-98 Hz)** | **1 to 7** | **3 patients without UPDRS** |
| 32 | Lofredi 2023<sup>47</sup> | 103 | n/a | **0.21** | **0.03** | **13-20** | **Off** | **UPDRS-III** | **No** | **yes (5-98 Hz)** | **1 to 7** | **3 patients without UPDRS scores** |
| 33 | Pardo-Valencia 2023<sup>48</sup> | 13 | 21 | 0.05 | n/a | 13-35 | Off-On | UPDRS-III | Yes | yes (460-490 Hz) | 3 to 5 |  |
| 34 | Pardo-Valencia 2023<sup>48</sup> | 13 | 21 | 0.1 | n/a | 13-35 | Off-On | UPDRS-III | No | yes (460-490 Hz) | 3 to 5 |  |
| 35 | Pardo-Valencia 2023<sup>48</sup> | 21 | 33 | 0.06 | n/a | 13-35 | Off | UPDRS-III | Yes | yes (460-490 Hz) | 3 to 5 | one-tailed Bayes factor 0.296 -> moderate evidence for absence of correlation |
| 36 | Pardo-Valencia 2023<sup>48</sup> | 21 | 33 | 0.17 | n/a | 13-35 | Off | UPDRS-III | No | yes (460-490 Hz) | 3 to 5 | one-tailed Bayes factor 0.550 -> anecdotal evidence for absence of correlation |
| 37 | Wiest 2023<sup>13</sup> | 14 | 28 | -0.04 | 0.85 | 13-35 | Off-On | Contra BR | Yes | yes (1-90 Hz) | 3 to 6 |  |
| 38 | Wilkins 2023<sup>49</sup> | 21 | 42 | LME | 1 | 13-30 | Off | Contra R | Yes | no | 3 to 70 months |  |
| 39 | Wilkins 2023<sup>49</sup> | 21 | 42 | LME | 1 | 13-30 | Off | Contra B | Yes | no | 3 to 70 months |  |
|  | **This study** | **99** | **178** | **0.25** | **0.01** | **13-20** | **Off-On** | **UPDRS-III** | **No** | **no** | **2.7 ± 1.8** |  |
|  | **This study** | **94** | **164** | **0.31** | **0.004** | **13-20** | **Off-On** | **UPDRS-III** | **Yes** | **no** | **2.7 ± 1.8** |  |
|  | **This study** | **99** | **178** | **0.31** | **0.002** | **13-20** | **Off-On** | **UPDRS-III** | **No** | **yes (5-95 Hz)** | **2.7 ± 1.8** |  |
|  | **This study** | **119** | **216** | **-0.04** | **0.69** | **13-20** | **Off** | **UPDRS-III** | **No** | **no** | **2.7 ± 1.8** |  |
|  | **This study** | **119** | **216** | **0.28** | **0.003** | **13-20** | **Off** | **UPDRS-III** | **No** | **yes (5-95 Hz)** | **2.7 ± 1.8** |  |
|  | **This study** | **119** | **216** | **0.24** | **0.01** | **13-20** | **Off** | **UPDRS-III** | **Yes** | **no** | **2.7 ± 1.8** |  |
| **Median** |  | **13** | **22** |  |  |  |  |  |  |  |  |  |

Significant positive correlations ( $p < 0.05$ ) are bold. B: bradykinesia, R: rigidity, Contra BR(T): contralateral bradykinesia-rigidity(-tremor) UPDRS subscore, LME: linear mixed effect model.

## 1 Multicenter reproducibility

In Fig. S2d-f, we averaged band powers of the left and right STNs. To account for the variability in analysis strategies used in previous studies, we performed the same correlations by sampling each STN (instead of each patient) and the corresponding contralateral UPDRS subscores for bradykinesia and rigidity (UPDRS-III items 22 to 26, Fig. S2c, h-k). On the single dataset level, we observed two significant negative correlations for the Oxford dataset (alpha-beta and beta) and a positive correlation for Düsseldorf2 and the low beta band. The pooled correlation coefficients were insignificant for the alpha-beta and beta bands (*ρ* _*α β*_ =0.03, *p* = 0.61 andρ _β_ = 0.09,*p* = 0.19) and significant for the low beta band (*ρ* _*L β*_*=* 0.17,*p* = 0.01). Note, however, that the estimated sample size to achieve proper replicability is *n* = 261 STNs. For this investigation, only *n* = 216 STNs were available.

We considered whether variability in beta peak frequencies across patients and STNs might affect correlations. Specifically, we investigated whether individual peak power, rather than power at fixed frequencies, might carry more meaningful pathological information. We, therefore, correlated the motor symptoms with the extracted maximum band powers.

However, the results shown in Fig. S2f were similar to those in panel b-with a negative correlation for the delta and theta bands (*p* < 0.05) and a positive correlation for the low beta band (*p* < 0.01). A scatter plot of the pooled low beta correlation from Fig. 2d is provided in Fig. S2g. Note that the mean low beta power correlation (ρ_mean(*L β*)_ = 0.26,p = 5e ^−3^) is similar to the maximum low beta power correlation (ρ_mean(*L β*)_ = 0.26,p = 0.01,) The panels d-g in Fig. S2 are also shown for the lateralized analysis in panels h-k. Instead of correlating baseline beta power with baseline UPDRS-III score in the Levodopa off condition, we also correlated the Levodopa modulation of the power in the beta band with the improvement of concomitant motor symptoms in Fig S3. The results for the baseline Levodopa on condition are shown in Fig. S4.

**Supplementary Fig. 2.**
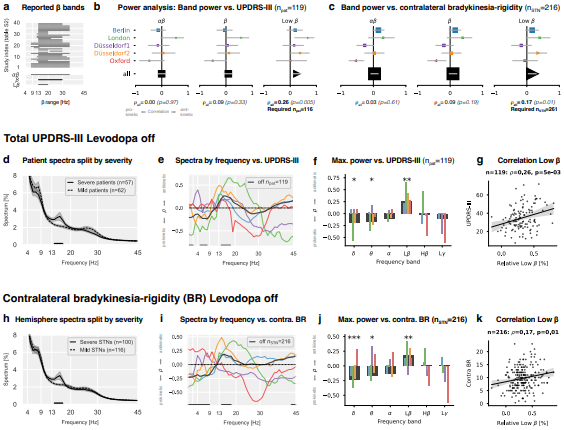
Association between relative spectral power and motor symptom severity for single and pooled datasets. **a**, Top: Beta band definitions used in previous studies (Table S2) correlating beta power with PD severity. Bottom: Frequency borders for the alpha-beta (8-35 Hz), beta (13-30 Hz), and low beta (13-20 Hz) bands used in this study. **b**, Correlations between average band power (left and right STNs averaged) and total UPDRS-III scores. X-axis: Spearman’s correlation coefficients, y-axis: datasets, horizontal lines: 95th percentile confidence intervals, symbol sizes represent the dataset sample sizes. Markers for non-significant correlations are displayed as squares and significant correlations as triangles. The pooled correlation coefficients and p-values are shown at the bottom. “Required n_pat_” indicates the sample size estimations to achieve a statistical power of 80% for the observed correlation coefficients. **c**, Same as **b**, treating each STN (left or right) as a single sample and correlating it with the contralateral bradykinesia-rigidity UPDRS score. **d**, Patient spectra split by their median UPDRS-III score. The horizontal line shows a 14-17 Hz cluster of significant difference. **e**, Correlation between power and UPDRS-III for each frequency bin of the power spectrum. Horizontal lines indicate frequencies with p-values < 0.05 for the pooled data. **f**, Correlations for maximum band power instead of mean band power. **g**, The scatter plot of the strongest pooled correlation from **b. h-k**, Same as **d-g;** however, without averaging left and right STNs, the contralateral bradykinesia-rigidity score instead of the total UPDRS-III score is used as a target.

**Supplementary Fig. 3.**
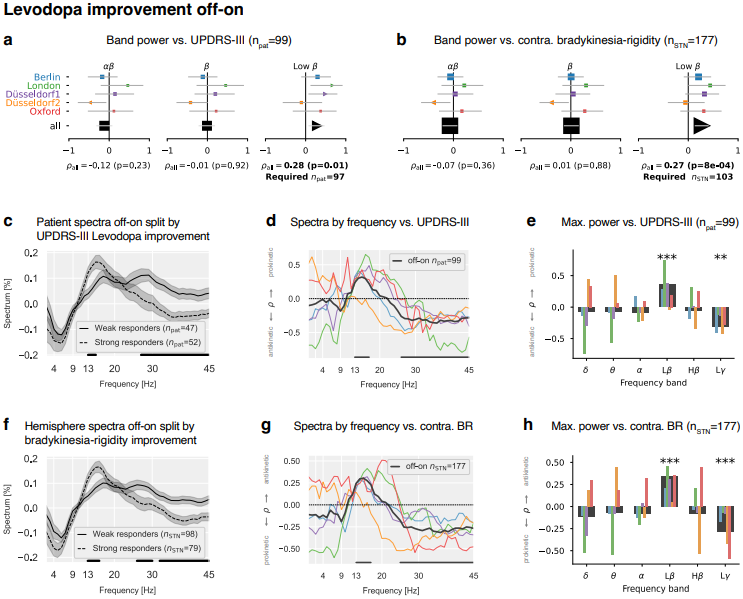
Same as **Fig. 3** for the Levodopa off-on improvement instead of the Levodopa off condition. The band power corresponds to the Levodopa-induced change of the power, and the UPDRS scores correspond to the Levodopa-induced symptom improvement.

**Supplementary Fig. 4.**
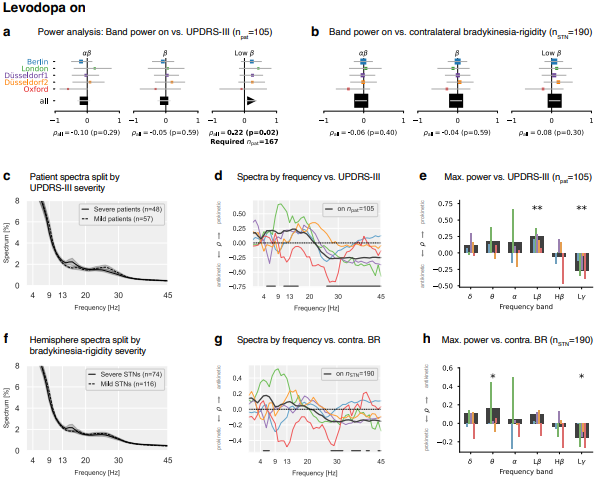
Same as **Fig. 3** for the Levodopa on condition.

## Stun effect

UPDRS assessment was performed only pre-surgically for Oxford and London and only post-surgically for Düsseldorf2, while for Berlin and Düsseldorf1, both assessment time points were available (Fig. 2a). We compared pre- and post-surgical UPDRS scores for Berlin and Düsseldorf1 and observed significantly lower scores for the off- and off-on condition (Fig. S5a), possibly indicating a stun effect^50^. Moreover, the difference between the pre- and post-UPDRS scores (post-pre) correlated negatively with the number of recovery days after the surgery for the off-and-on condition, indicating that the stun effect was more pronounced for fewer days between surgery and measurement (Fig. S5b).

We further investigated the impact of the stun effect on the LFP power using the Berlin, Düsseldorf1, and Oxford datasets. London and Düsseldorf2 were excluded because, in those datasets, all recordings were performed on a fixed number of days between surgery and recording. Correlating the number of days between surgery and recording with LFP power revealed that low-frequency power (delta, theta, alpha) decreased with more days since surgery. In contrast, low beta, high beta, and low gamma power increased (Fig. S5c). This indicates that beta power is suppressed after the surgery and rebounds afterward.

Despite these important factors impacting both the STN spectrum and the UPDRS assessment, the correlations for the pre- and post-operative UPDRS scores were similar (*ρ*_*L β*_ off pre = 0.24, *ρ*_*L β*_ off post = 0.21, *ρ*_*L β*_ off-on pre = 0.46, *ρ*_*Lβ*_ off-on post = 0.42,Fig. S5d), in line with Pardo-Valencia^48^. These findings suggest that the most realistic UPDRS symptom assessment (unaffected by the stun effect) is achieved pre-operatively or many days after the surgery. However, the stun effect appears to reduce beta power and UPDRS scores similarly, therefore having no strong impact on the reported correlations. Finally, no surgical microelectrode recordings were performed for the London and Oxford datasets (Table 3), possibly reducing the strength of the stun effect^51^.

**Supplementary Fig. 5.**
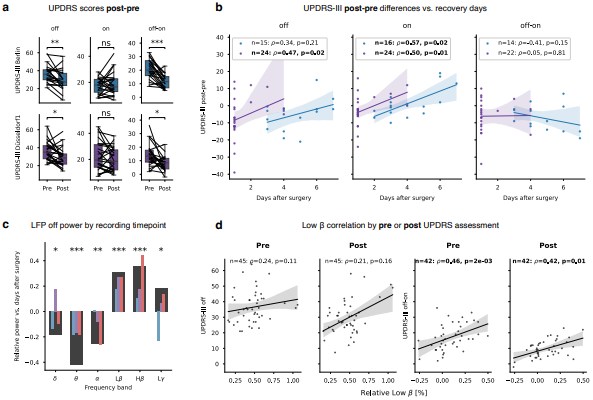
Stun effect. **a**, Comparison of pre- and postoperative UPDRS-III scores. Pre- and post-operative scores were available for patients from Berlin and Düsseldorf1 (c.f. Fig. 2a). **b**, The difference in UPDRS scores (post-pre) decreases with increasing recovery days after the surgery. Same colors as in **a. c**, Delta, theta, and alpha decrease during recovery. Low beta, high beta, and low gamma power increase during recovery. Same colors as in **a**. Red: Oxford. Black: All. We excluded London and Düsseldorf2 from this analysis because all patients’ recovery days were equal, prohibiting a correlation analysis (c.f. Fig. 2a). **d**, Comparison of correlating beta power with the pre-vs. post-operative UPDRS scores for Berlin and Düsseldorf1 pooled.

## 3 Within-patient correlations

**Supplementary Fig. 6.**
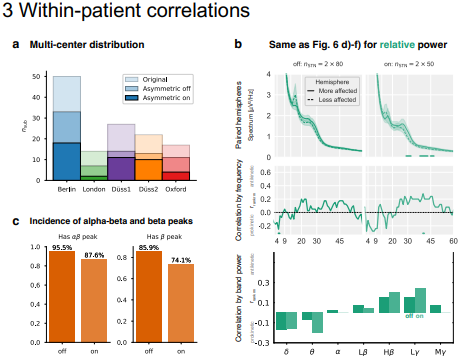
Supplementary information for within-patient investigation in Figs. 6-7. **a**, Within-patient analysis inclusion by dataset and Levodopa condition. **b**, Same as **Fig. 6 d-f** for relative power. **c**, *specparam*’s probability of fitting a peak in a given frequency band. Same as **Fig. 7b** for alpha-beta and beta. Only when considering the full alpha-beta range (8-35 Hz) did at least 88% of STNs have a peak, as previously shown^52–54^.

**Supplementary Fig. 7.**
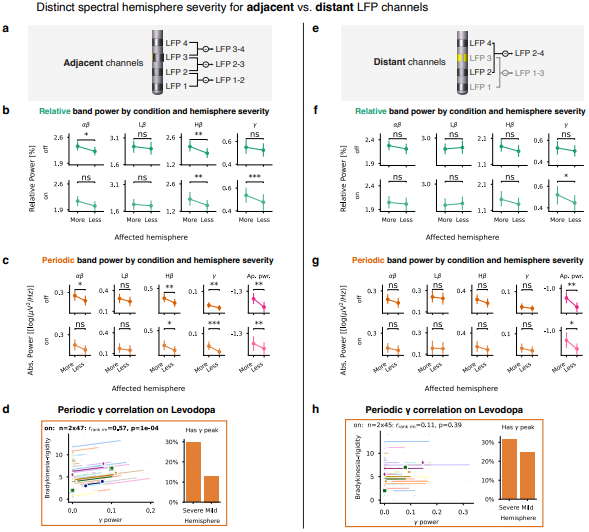
Distinct spectral hemisphere severity for adjacent vs. distant LFP channels. **a**, Visualization of adjacent bipolar LFP montage. **b**, Wilcoxon signed-rank test for relative band power comparing the more and less affected hemispheres in the Levodopa off- and on-state. **c**, Same as **b** for absolute periodic power. **d**, Left: Repeated measures correlation for periodic gamma oscillations in the Levodopa on-state. Four highlighted subjects are shown in **Supplementary Fig. 9d**. Right: *specparam*’s probability of fitting gamma peaks by hemisphere. **e-h**, Same as **a-d** for distant LFP channels recommended for aDBS. Note that alpha-beta and gamma are only elevated for the more affected hemisphere in the Levodopa off-or on-state, respectively, and they are not robust with regard to the spatial specificity of the bipolar montage. In contrast, aperiodic broadband power is elevated in the more affected hemisphere independent of Levodopa medication and channel choice.

DBS electrodes enable LFP sensing, usually via a bipolar reference montage. Some studies focus on adjacent DBS channels (1-2, 2-3, or 3-4, Fig. S7a), whereas others use distant channels (1-3 or 2-4, Fig. S7e). Adjacent channels better capture local dynamics, whereas distant channels are spatially less specific. When analyzing adjacent bipolar channels, we observe a strong within-patient correlation (*r*_*rank rm*_ = 0.57,*p* = 1e^−4^) between bradykinesia-rigidity symptoms and periodic gamma oscillations in the Levodopa on-state (Fig. S8b).

These STN gamma oscillations are highlighted in yellow in Fig. S8d and were reported before^55,56^. However, their local specificity limits their potential as a reliable aDBS biomarker. An optimal aDBS biomarker should be detectable from distant LFP channels, such as 1-3 or 2-4, which are adjacent to the central stimulation contacts (2 or 3). For distant channels (1-3 or 2-4), only absolute total mid gamma power and aperiodic broadband power correlated significantly with patients in the Levodopa off- and on-state.

**Supplementary Fig. 8.**
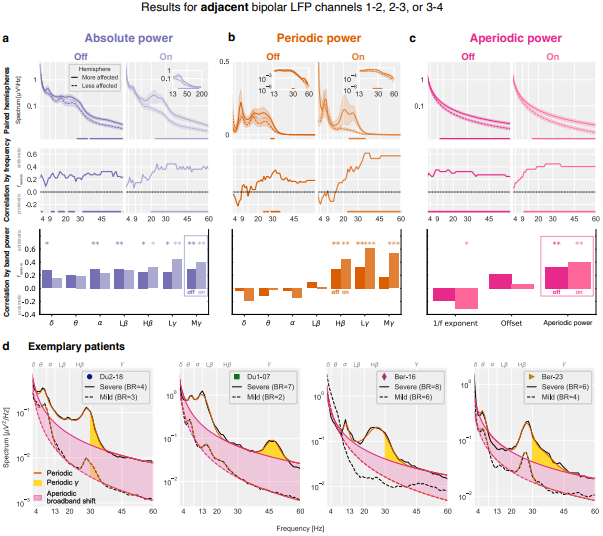
Periodic gamma oscillations correlate with motor symptom severity within patients on Levodopa. Same as Figs. 6-7 for adjacent instead of distant bipolar LFP channels. **a**, Top: Within-patients paired cluster-based permutation tests for the more and less affected hemisphere. Middle: Repeated measures rank correlation for each frequency bin of the power spectrum and the contralateral bradykinesia-rigidity score. Bottom: Repeated measures rank correlation for band powers. **b-c**, Same as **a** for periodic and aperiodic power. **d**, Exemplary patients. The symbols of each patient correspond to the symbols in the repeated measures scatter plots in **Fig. S7d** and **S7h**. The gamma oscillations often overlap with high beta oscillations (Du2-18, Ber-16, Ber-23) but correlate stronger with symptoms than high beta power. Note that the periodic gamma oscillation in patient Du1-07 is more prominent in the adjacent bipolar montage compared to the distant bipolar montage shown in **Fig. 7**. This suggests that periodic gamma oscillations carry pathological information but are spatially narrowly localized, hindering their utility as aDBS biomarker.

